# Hippocampal-hypothalamic circuit controls context-dependent innate defensive responses

**DOI:** 10.1101/2021.08.06.455454

**Authors:** Jee Yoon Bang, Julia Sunstrum, Danielle Garand, Gustavo Morrone Parfitt, Melanie Woodin, Wataru Inoue, Jun Chul Kim

**Author notes:** **Corresponding author:** Dr. Jun Chul Kim, University of Toronto, Department of Psychology, 100 St. George Street, Toronto, ON, M5S 3G3, Canada, (lab). **Funding source** Ontario Graduate Scholarship (OGS) [JYB]. BrainsCAN Studentship and NSERC-CGSD [JS]. NSERC Discovery MOP 506730 and CIHR MOP 507489 [JCK]. NSERC Discovery MOP 482622 [MW]. CIHR MOP 148707 [WI].

## Abstract

Preys use their memory - where they sensed a predatory threat and whether a safe shelter is nearby - to dynamically control their survival instinct to avoid harm and reach safety. However, it remains unknown which brain regions are involved, and how such top-down control of innate behaviour is implemented at the circuit level. Here, we show that the anterior hypothalamic nucleus (AHN) is best positioned to perform this task as an exclusive target of the hippocampus (HPC) within the medial hypothalamic defense system. Selective optogenetic stimulation and inhibition of hippocampal inputs to the AHN revealed that the HPC→AHN pathway not only mediates the contextual memory of predator threats but also controls the goal-directed escape by transmitting information about the surrounding environment. These results reveal a new mechanism for experience-dependent, top-down control of innate defensive behaviours.

## Main

Manoeuvring through a rapidly changing environment while avoiding the threat of predation is essential for the survival and reproduction of all species^1^. This requires abilities to perceive the magnitude of predator threats (i.e., stimulus detection and integration), initiate defensive responses such as escape flight or freezing (i.e., defensive motor actions), and in parallel, remember the area where the predator appeared (i.e., memorization) so that the possibility of re-encountering the same threat can be avoided^2, 3^. Upon detecting predatory threats, prey animals also select the most successful defense strategy based on their knowledge of the surrounding environment such as the presence of nearby food and the availability of a safe shelter^4, 5^. For example, when there is no safe shelter, rodents select freezing over escape flight to avoid being detected by predators. Once they learn about the existence of a safe shelter, however, defense strategy quickly switches to escape-running toward the shelter^6, 7^. Thus, defensive response to predatory threats is not simple stimulus-response, but a flexible, cognitive process that utilizes the knowledge of prior experiences and environments^4, 6, 8^.

Innate defensive behaviours are generated by the medial hypothalamic defensive system^9^, consisting of the anterior hypothalamic nucleus (AHN)^10, 11^, the dorsomedial and central region of the ventromedial hypothalamus (VMHdm/c)^10, 12–14^ and the dorsal premammillary nucleus (PMD)^9, 15^. These three distinct nuclei are densely interconnected and become highly active upon predator exposure^16–19^ to control motor outputs at the level of periaqueductal gray (PAG)^7, 12, 20, 21^. In both rodents and non-human primates, direct stimulation of the medial hypothalamic defensive system evokes strong defensive responses, such as escape flight, freezing, sympathetic activation, and panic, while its inhibition reduces defensive responses to predator threats^22–25^.

How are then the hard-wired defensive responses flexibly controlled by animals’ memory and knowledge of the environment? While the medial hypothalamus defense system has been extensively studied, it remains unknown how information about threat-associated context and spatial environment is implemented at the circuit level during the innate defensive response to predator threats. It is well-established that the environmental context of a salient event is first encoded within the hippocampus (HPC) as the collective activity of place cells and time cells ^2, 3, 26–30^. Later during memory recall, the contextual information serves as a potent retrieval cue by reinstating patterns of brain activity observed during the original experience.^31–33^

Given the critical role of the hippocampus in encoding contextual memory, we hypothesized that hippocampal inputs to the medial hypothalamic defensive system may control innate defensive responses based on the animals’ knowledge of the surrounding environment. Using a combination of anterograde tracing and electrophysiological recording, we first found that the hippocampus innervates almost exclusively the AHN (i.e., HPC→AHN pathway), but not the PMD or VMHdm/c. Subsequent optogenetic activation and inhibition experiments showed that the HPC→AHN pathway not only mediates the contextual memory of predator threats but also controls the goal-directed escape by transmitting information about the surrounding environment.

## Results

### AHN stimulation evokes escape responses

To examine the behavioural consequences of anterior hypothalamic nucleus (AHN) activation, we transduced neurons in the AHN by bilateral injection of adeno-associated viral vector (AAV) with human synapsin promoter (hSyn) carrying channelrhodopsin-2 (AAV-hSyn-ChR2-GFP) or GFP (AAV-hSyn-GFP) (Fig. 1a,b). The location of viral transduction and optic fiber placement were confirmed to be in the central and caudal regions of AHN with minimal spread to neighbouring hypothalamic areas (Extended Data Fig. 1a,b). We first examined the behavioural effects of low and high frequency (6 Hz and 20 Hz) stimulation and found that the high frequency stimulation generated robust escape behaviours in the absence of any overt predator threat (Extended Data Fig. 2). To systematically investigate the behavioural effects of AHN stimulation, we optogenetically stimulated the AHN in three different escape conditions with varying degrees of difficulty (Fig. 1c,d): 1) an open field arena with short transparent walls (condition 1, easy), tall opaque walls (condition 2, hard), and physical restraint tube (condition 3, impossible). In condition 1, AHN stimulation induced bursts of running (> 0.3 m/s) with a short latency (5 ± 1.29 s) (Supplementary video 1). After bouts of running, AHN-ChR2 mice, but not GFP-control mice, initiated multiple escape jumps which resulted in 5 of 6 AHN-ChR2 mice escaping the test arena. We quantified the light-induced behavioural effect as a normalized difference between baseline epoch (OFF, 2 min) and stimulation epoch (ON, 2 min) and found that AHN-ChR2 mice had significant increases in the speed of locomotion, freezing, and jumping (Fig. 1e-g) compared to GFP controls. In condition 2 (hard), no AHN-ChR2 mice escaped the test arena, but escape attempts were maintained with increased running, freezing, and jumping compared to GFP controls (Fig. 1h-j, Supplementary video 2). In condition 3, animals were physically restrained and received AHN stimulation (10s ON, 10 s OFF) for 30 minutes, during which escape struggle movements were visually inspected and monitored using a collar sensor with a pulse oximeter. Despite limited mobility and the long duration of physical restraint, AHN-ChR2 mice, but not GFP controls, displayed persistent escape-struggle movements throughout the test. Thus, our data demonstrate that AHN activity is sufficient to evoke escape-associated behavioural responses in the absence of overt predator cues.

**Figure 1.**
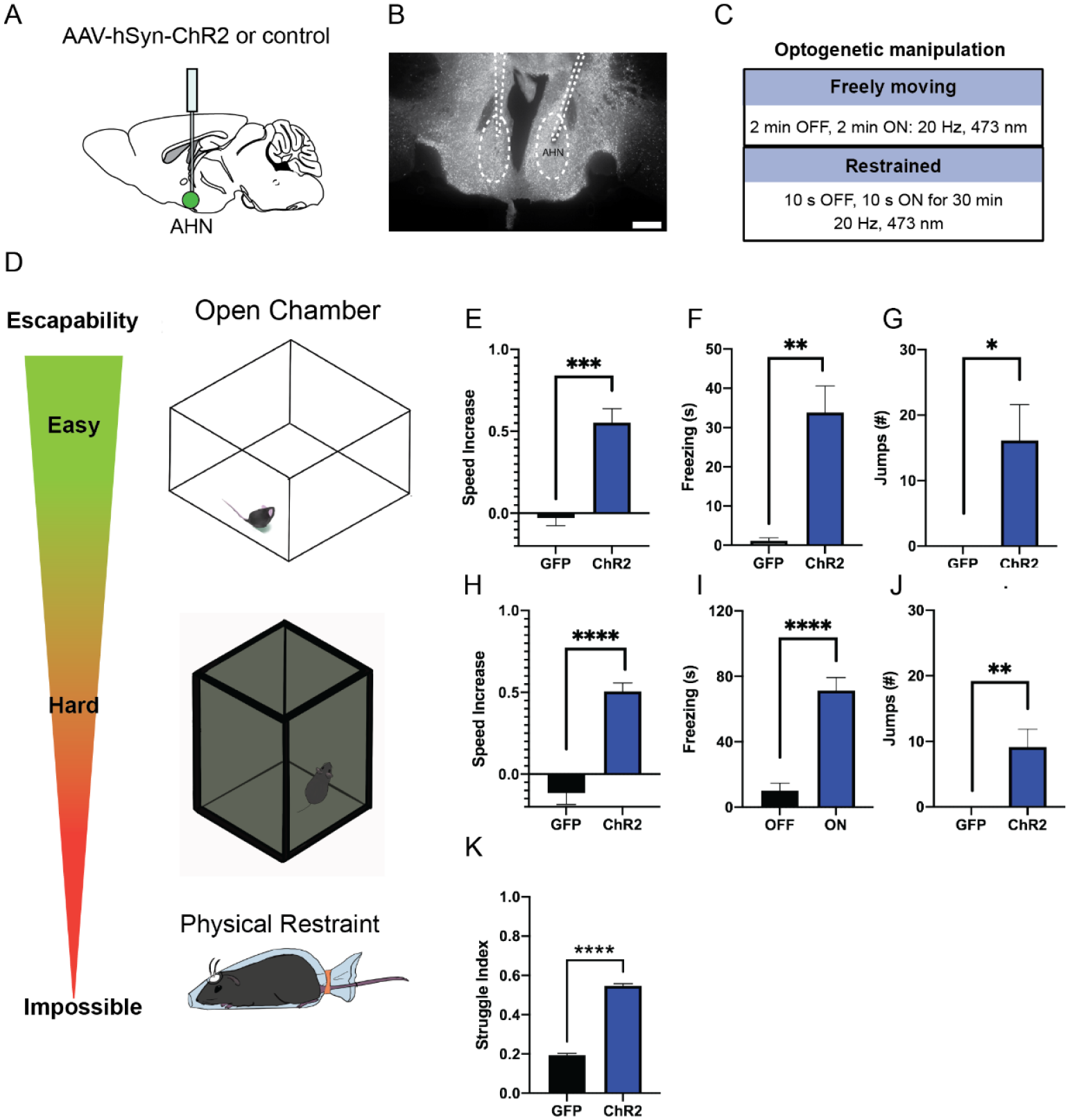
AHN stimulation induces escape-associated behaviours. **a**, Schematic illustration of optogenetic activation in the AHN (green circle depicts the AAV infusion). **b**, An example of histological confirmation showing the expression of ChR2 and placement of optic fiber in the AHN. **c**, Schematic describing optogenetic stimulation paradigm. **d**, Three different escape conditions where the effects of AHN stimulation was examined. Top: open field arena with short transparent walls (condition 1, easy). Middle: tall opaque walls (condition 2, hard). Bottom: physical restraint tube (condition 3, impossible). **e,** Condition 1: speed increase from the light OFF epoch to ON epoch (unpaired t-test, two-tailed, t=5.209, df=11, *\*\*\*p=0.0003*). **f**, Condition 1: freezing time during the light ON epoch (GFP N=7, ChR2 N=6, unpaired t-test, two-tailed, t=4.269, df=11, *\*\*p=0.0013*). **g**, Condition 1: number of jumps during the light ON epoch (unpaired t-test, two-tailed, t=2.308, df=11, *\*p=0.0414*). **h**, Condition 2: speed increase from the light OFF epoch to ON epoch (GFP N=7, ChR2 N=6, unpaired t-test, two-tailed, t=6.204, df=11, *\*\*\*\*p<0.0001*). **i**, Condition 2: freezing time during the light ON epoch (GFP N=7, ChR2 N=6, unpaired t-test, two-tailed, t=6.695, df=11, *\*\*\*\*p<0.0001*). **j**, Condition 2: number of jumps during the light ON epoch (GFP N=7, ChR2 N=6, unpaired t-test, two-tailed, t=3.796, df=11, *\*\*p=0.003*). **k**, Condition 3: struggle movement during the 30 minutes of physical restraint (GFP N=4, ChR2 N=6, unpaired t-test, two-tailed, t=24.74, df=382, \*\*\*\**p<0.0001*). All results reported are mean ± s.e.m. \**p < 0.05, **p < 0.01, ***p < 0.001, ****p<0.0001*. Scale bar= 200µm

### AHN activation carries negative valence and induces conditioned avoidance

Fear of predators is an aversive emotional state that elicits defensive behaviours such as freezing and escape flight^16, 34, 35^. Therefore, we probed the emotional valence of AHN activation in a close loop real-time place avoidance assay (RTPA) (Fig. 2a,b). During a 5 min habituation, mice were allowed to explore two distinct chambers, and a preferred chamber was selected as the photostimulation chamber (Fig. 2c,d). During a subsequent 20 min RTPA test, mice explored two chambers and received AHN stimulation at either low or high frequency (6 or 20 Hz) only in the photostimulation chamber (Fig. 2b). All AHN-ChR2 mice exhibited dramatic flight responses upon AHN activation, immediately leaving and avoiding the photostimulation chamber (Fig. 2e, Supplementary Video 3). While both 6 and 20 Hz stimulation induced significant avoidance of the photostimulation chamber in ChR2 animals, there was a frequency-dependent magnitude of aversion. The 20 Hz stimulation induced a greater mean aversion index (- 0.89) than the 6 Hz stimulation (−0.63) (Fig. 2f) with no difference in total distance travelled during the test (Fig. 2g). Since the AHN photostimulation was paired with a distinct context, we next asked whether the aversion evoked by AHN activity is sufficient to induce conditioned place avoidance (CPA). A day after the real-time place aversion test, mice were placed back in the middle of the two chambers, but without photostimulation, to determine their conditioned place aversion. Most AHN-ChR2 mice immediately turned away from the photostimulation chamber and exhibited investigatory behaviours towards the entrance of photostimulation chamber (Supplementary Video 3). Both 6 and 20 Hz stimulation produced significant conditioned place aversions with the 20 Hz stimulation inducing a greater mean aversion index (- 0.53) than the 6 Hz stimulation (- 0.20). Together, our data demonstrate that AHN activity carries negative emotional valence and can serve as a stimulus for the formation of a conditioned place aversion memory.

**Figure 2.**
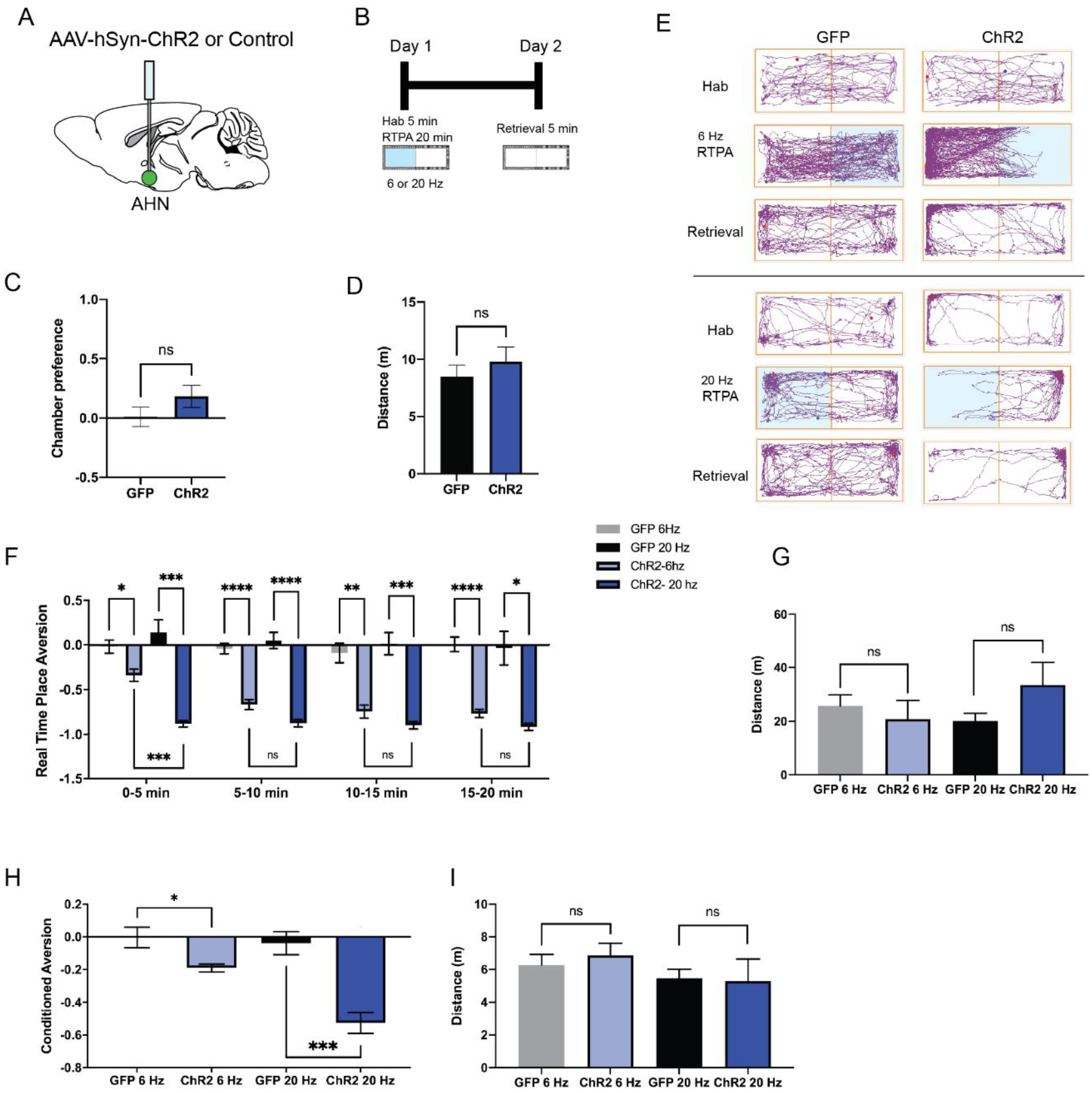
AHN stimulation is aversive and induces conditioned place aversion. **a**, Schematic illustration of optogenetic activation in the AHN (green circle depicts the AAV infusion). **b**, Schematic describing the RTPA and CPA test paradigm: day 1 consisting of habituation and real-time place preference (20 min) and day 2 for testing conditioned place preference (5 min). **c**, Chamber preference during habituation (GFP N= 14, ChR2 N=12, unpaired t-test, t= 1.390, df= 24, *p=0.1772,* NS). **d**, Distance travelled during habituation (GFP N= 14, ChR2 N=12, unpaired t-test, t=0.8396, df=24, *p=0.41*, NS). **e**, Representative locomotion trajectory for a GFP control animal (left column) and a ChR2-expressing animal (right column) during habituation (hab), 6 Hz or 20 Hz real-time stimulation (6 Hz RTPA, 20 Hz RTPA), and conditioned place aversion test (Retrieval). Light-coupled chambers are shown in blue. **f**, Real time place aversion monitored across 20 minute test (GFP N=7, ChR2 N=6). GFP 6 Hz vs. ChR2 6 Hz (2-WAY RM ANOVA, time x treatment F(3,33)=3.965, *\*p=0.016*). GFP 20 Hz vs. ChR2 20 Hz ( 2-WAY RM ANOVA, time x treatment, F(3,33)=0.6059, *p=0.6158*). GFP 6 Hz vs. GFP 20 Hz (2-WAY RM ANOVA, time x frequency, F(2.071, 12.42)=1.076*, p=0.3730*, NS). ChR2 6 Hz vs. ChR2 20 Hz (2-WAY RM ANOVA, time x frequency, F(1.514, 7.274)= 11.05, \**p=0.0223*). **g**, Distance travelled during 6 Hz and 20 Hz real-time stimulation. GFP 6 Hz vs. ChR2 6 Hz (unpaired t-test, t=0.6245, df=11, *p=0.545*). GFP 20Hz vs. ChR2 20 Hz (unpaired t-test, t=1.612, df=11, *p=0.1352*). **h**, Conditioned aversion memory tested 24-hours after real time place aversion tests. GFP 6 Hz vs. ChR2 6 Hz (unpaired t-test, t= 2.581, df=11, \**p=0.0256*). GFP 20 Hz vs. ChR2 20 Hz (unpaired t-test, t=5.044, df=11, \*\*\**p=0.0004*). ChR2 6 Hz vs. ChR2 20 Hz (paired t=test, t=7.642, df=5, \*\*\**p=0.0006*). GFP 6 Hz vs. GFP 20 Hz (paired t-test, t= 0.4013, df= 6, *p=0.7021*). **i**, Distance travelled during the conditioned place aversion test. GFP 6 Hz vs. ChR2 6 Hz (unpaired t-test, t=0.6084, df=11, *p=0.5553*). GFP 20 Hz vs. ChR2 20 Hz (unpaired t-test, t=0.9016, t=0.1265, df=11, *p=0.9016*). All results reported are mean ± s.e.m. \**p < 0.05, **p < 0.01, ***p < 0.001, ****p<0.0001*.

### AHN receives direct glutamatergic inputs from the hippocampus

Next, we investigated the distribution of the hippocampal fiber afferents to the hypothalamus. To this end, we performed anterograde tracing from the hippocampus using virally delivered-ChR2 (AAV-hSyn-ChR2-GFP) as an anterograde tracer (Fig. 3a, Extended Data Fig. 3a). HPC infusions led to the expression of ChR2-GFP in the ventral two-thirds of the HPC, with minimal spread into adjacent cortical structures such as the entorhinal cortex which does not project to the AHN (Extended Data Fig. 3b). Consistent with previous reports, GFP-positive axon terminals were detected in the known targets of HPC, including the amygdala^36, 40^, lateral septum^36–40^, nucleus accumbens^40, 41^ and prefrontal cortex^40–42^ (data not shown). Within the hypothalamus, HPC axon terminals were found most abundantly in the AHN based on a normalized measure of GFP fluorescence intensity (Fig. 3b,c). In stark contrast, the VMHdm/c and PMD, the other two main components of the medial hypothalamic defense system were almost excluded from the HPC innervation. Furthermore, the hippocampal innervation of the AHN showed an overall bias against other medial hypothalamic nuclei implicated in stress-induced corticosterone release (PVN) and social aggression (VMHvl) (Fig. 3b,c). This data indicates that the AHN is the primary entry point for HPC inputs to the medial hypothalamic defense system.

**Figure 3.**
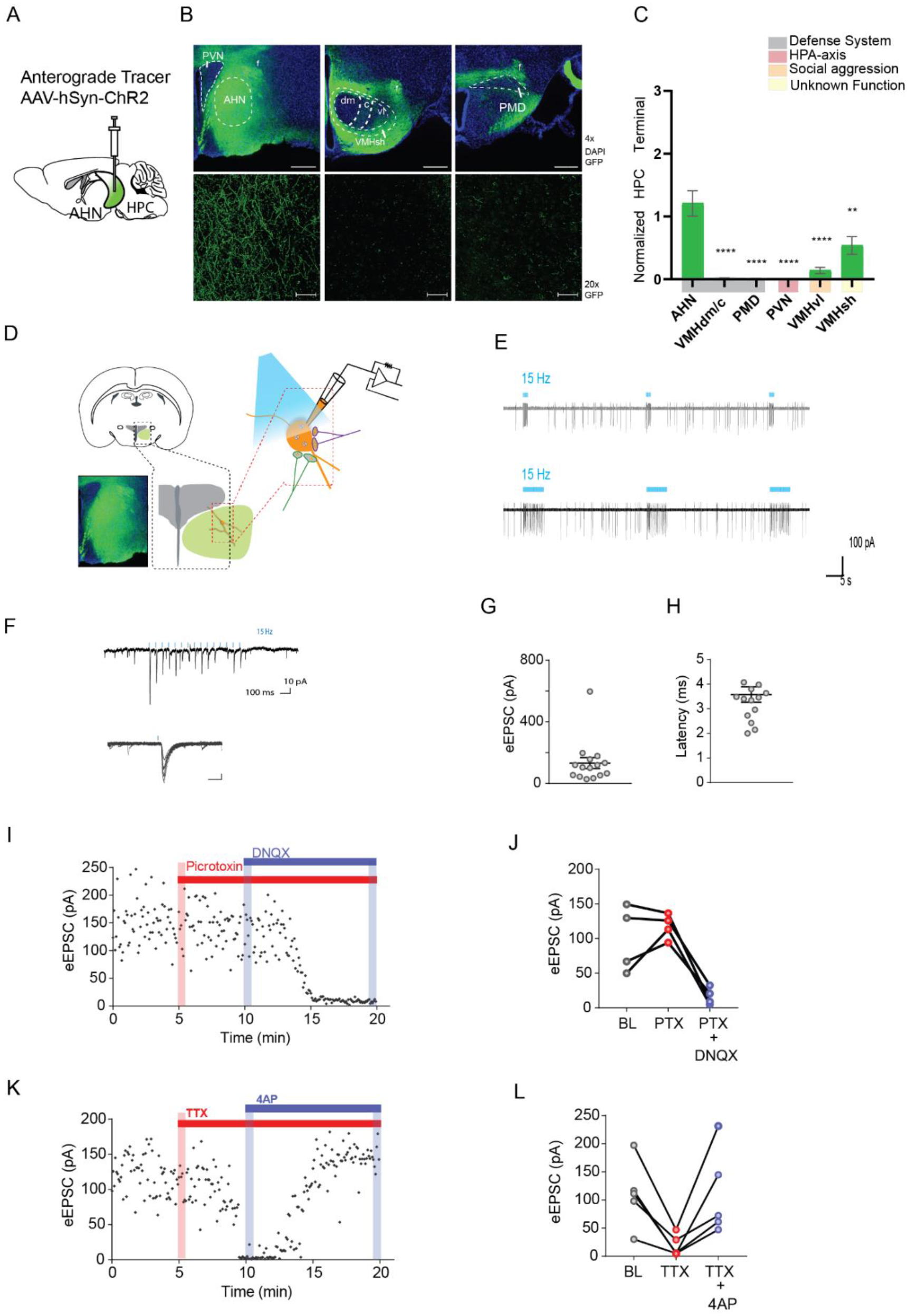
Hippocampus sends monosynaptic excitatory inputs to the anterior hypothalamic nucleus. **a**, Schematic illustration of anterograde tracing experiment. **b**, HPC terminals (green) in the hypothalamus, including anterior hypothalamic nucleus (AHN), dorsomedial and central regions of ventromedial hypothalamus (VMHdm/c), premammillary dorsal nucleus (PMD), paraventricular nucleus (PVN), ventrolateral region of ventromedial hypothalamus (VMHvl), shell of ventromedial hypothalamus (VMHsh). DAPI staining (blue). **c**, Quantification of HPC terminal intensity (N=2, One-Way ANOVA, multiple comparison Dunnett’s test, *\*\*\*\*p<0.0001, **p<0.01*). **d**, Schematic illustration for patch clamp recordings of AHN neurons in coronal brain slices that express ChR2 in HPC terminals. **e,** Examples of cell attach recordings. Illumination of blue light (480 nm, 5 ms pulse at 15 Hz) triggered firing of AHN neurons. **f**, Examples of whole-cell voltage-clamp recordings of AHN neurons. Blue light illumination (5 ms) evoked inward current. **g**,**h**, Summary of light evoked EPSCs (**g**) amplitude and latency (**h**). **i**, Light-evoked EPSCs persisted in the presence of GABA A receptor antagonist picrotoxin (PTX, 100 µM) and eliminated by AMPA/kainite receptor antagonist DNQX (20 µM). **j**, Summary of eEPSC change after PTX and DNQX application. **k**, Light-evoked EPSCs were eliminated by TTX (0.5 µM) and then recovered by a low dose 4-AP (100 µM). **l**, Summary of eEPSC changes after TTX and 4-AP application. All results reported are mean ± s.e.m. *\*p < 0.05, **p < 0.01, ***p < 0.001, and ****p<0.0001.* Scale bar= 100µm

To further validate direct hippocampal inputs arriving at the AHN and determine their electrophysiological properties, we carried out cell-attached and whole cell patch-clamp recordings from AHN cells in acute brain slices containing ChR2-expressing HPC axon terminals (Fig. 3d). In the cell-attached voltage-clamp mode, photostimulation of HPC terminals (473 nm, 5 ms pulses at 15 Hz) triggered robust action potential firings of AHN cells (Fig. 3e). In the whole-cell voltage clamp mode, photostimulation induced short-latency (average latency 3.6 ms) excitatory postsynaptic currents (EPSCs) (average amplitude 132 pA) (Fig. 3f-h). Light-evoked EPSCs in the AHN were not affected by GABAA receptor antagonist picrotoxin (PTX, 100 µM) but eliminated by AMPA/kainite receptor antagonist DNQX (10 µM), indicating that HPC input to the AHN is glutamatergic in nature (Fig. 3i,j). To isolate monosynaptic inputs from ChR2-expressing HPC axons, we sequentially added tetrodotoxin (TTX, 1 μM) and 4-aminopyridine (4-AP, 100 μM) to the ACSF. The previously observed light-evoked EPSCs were eliminated by TTX but recovered after the application of 4-AP, lending further support that monosynaptic transmission was triggered by direct ChR2-mediated depolarization of HPC terminal boutons^43^ (Fig. 3k,l).

As AHN is heavily populated by GABAergic cells, we next investigated whether the direct HPC innervation of AHN is biased toward GABA cells. We repeated the current clamp recording experiments with AHN slices from double transgenic reporter mice (RC::Frepe, Dlx5/6-FLPe, Extended Data Fig. 4a) in which forebrain GABA cells are specifically labelled with red fluorescent protein, mCherry. We observed mCherry labelled GABA cells in the AHN but not in VMHdm/c and PMD (Extended Data Fig. 4b, bottom row). As expected, photostimulation evoked action potential spikes in mCherry-positive AHN GABA cells. However, we did not find any significant difference in the number of photostimulation-induced spikes between mCherry-positive and mCherry-negative cells, indicating that HPC axon terminals synapse on both GABA and glutamatergic cells in the AHN (Extended Data Fig. 4c-i).

Together, our anterograde tracing and electrophysiological recording demonstrate that the AHN receives direct monosynaptic excitatory inputs from the HPC. These findings also suggest that the AHN plays a specialized role in the medial hypothalamic defensive system, different from other major components, namely the PMD and VMHdm/c.

### Activation of HPC→AHN pathway induces escape-associated locomotion

The hippocampus sends direct monosynaptic inputs to the AHN, but their behavioural function remains unknown. Thus, we examined if activating HPC→AHN pathway would induce the same behavioural responses seen in the direct AHN soma activation. The HPC was virally transduced with AAV-hSyn-ChR2-GFP, and optic fibers were bilaterally implanted at the AHN to illuminate HPC axon terminals (Fig. 4a,b). The viral transduction was confirmed to include all hippocampal presynaptic sources of the AHN along the entire dorsoventral axis of the hippocampal formation (dSUB, dCA1, vCA1, vSUB) (Extended Data Fig. 1c,d). Light induced-behavioural changes were then monitored during low or high frequency (6 or 20 Hz) stimulation of HPC→AHN pathway in an open field arena and compared between ChR2 and GFP control mice (Fig. 4c,d, Extended Data Fig. 5). To our surprise, the pathway activation did not elicit robust escape jumping or freezing observed in the direct AHN activation. Instead, it produced light-synched, reversible increases in running bouts and speed (Fig. 4e, Extended Data Fig. 5c). We also examined light-induced changes in consummatory (rearing, grooming) behaviours (Fig. 4g,h, Extended Data Fig. 5), and found that only grooming was decreased during the 6 Hz light stimulation (Fig. 4f-h, Extended Data Fig. 5f). To corroborate the effects of activating HPC→AHN pathway on escape-associated locomotion, we repeated the same pathway activation during physical restraint condition. The delivery of bursts of light pulses (20 Hz, 10s ON, 10s OFF) for 30 minutes significantly increased escape-associated struggle movements in ChR2 mice compared to controls (Fig. 4i), consistent with the effect of direct AHN soma activation. Thus, our data suggests that HPC→AHN pathway activity promotes escape responses by inducing locomotion.

**Figure 4.**
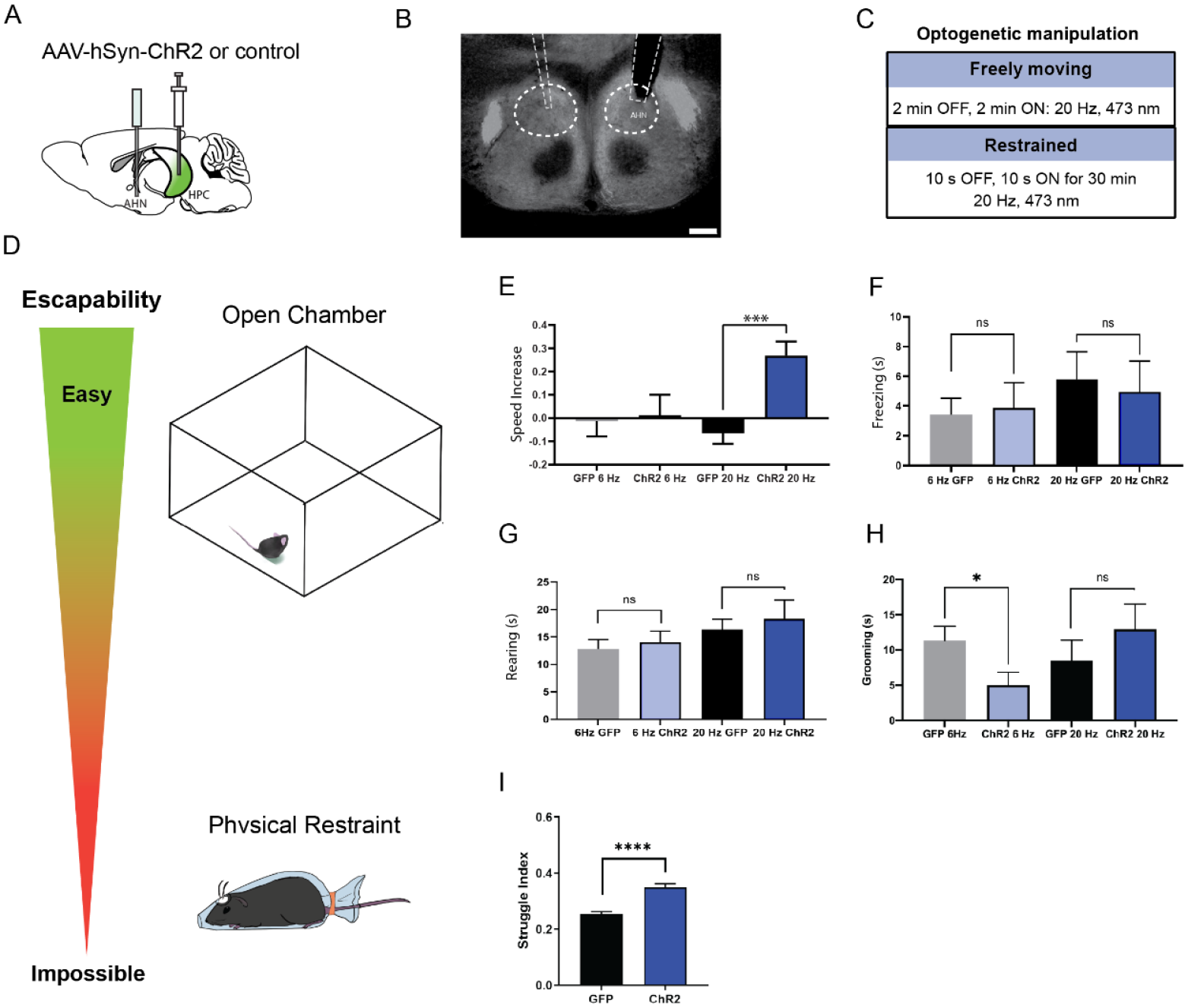
HPC→AHN pathway activation induces escape-associated locomotion. **a,** Schematic illustration of optogenetic activation of hippocampal terminals in the AHN (GFP N=10, ChR2 N=10). **b**, An example of histological confirmation showing the expression of HPC terminals and placement of optic fibers in the AHN. **c**, Schematic describing optogenetic stimulation paradigm. **d**, Two different escape conditions where the effects of HPC terminal stimulation was examined. Top: open field arena with short transparent walls (condition 1, easy). Bottom: physical restraint tube (condition 2, impossible). **e**, Condition 1: speed increase from the light OFF epoch to ON epoch. GFP 6 Hz vs. ChR2 6 Hz (unpaired t-test, two-tailed, t=0.2281, df= 18, *p= 0.8222*, NS). GFP 20 Hz vs. ChR2 20 Hz (unpaired t-test, two-tailed, t=4.366, df=18, *\*\*\*p=0.0004*). **f**, Condition 1: freezing time during the light ON epoch. 6Hz GFP vs. 6Hz ChR2 (unpaired t-test, two-tailed, t=0.2035, df=18, *p=0.8271*, NS). 20Hz GFP vs. 20 Hz ChR2 (unpaired t-test, two-tailed, t= 0.1323, df=18*, p=0.8962*, NS). **g**, Condition 1: rearing time during the light ON epoch. GFP 6 Hz vs. ChR2 6 Hz (unpaired t-test, two-tailed, t=0.0308, df=18, p=0.9758, NS). GFP 20 Hz. vs. ChR2 20 Hz (unpaired t-test, two-tailed, t=, df=18, *p=0.7794,* NS). **h**, Condition 1: grooming time during the light ON epoch. GFP 6 Hz vs. ChR2 6 Hz (unpaired t-test, two-tailed, t=2.244 df=18, *\*p=0.0376*). GFP 20 Hz vs. ChR2 20 Hz (unpaired t-test, two-tailed, t=0.6362, df=18*, p=0.5327*, NS). **i**, Condition 2: struggle movement during the 30 minutes of physical restraint (unpaired t-test, two-tailed, t=6.685, df=424 *\*\*\*\*p<0.0001*). All results reported are mean ± s.e.m. *\*p < 0.05, **p < 0.01, ***p < 0.001, ****p<0.0001*. Scale bar= 200µm

### HPC→AHN pathway activation is aversive and instructs learning of a conditioned place aversion

To evaluate whether the HPC→AHN pathway activity is intrinsically aversive and sufficient to induce a conditioned place aversion, we used the RTPA and CPA paradigms (Fig. 5a,b). During the habituation, mice did not show any significant preference to either chamber, and distance travelled did not differ (Fig. 5c,d). During the subsequent RTPA task, ChR2 mice gradually developed an avoidance to a chamber paired with light stimulation at both 6 Hz and 20 Hz frequency (Fig. 5e,f, Supplementary video 4, 5). Although there was a trend of increased locomotion with stimulation at 20 Hz frequency, total distance travelled did not differ compared to the controls (Fig. 5g). One day after the RTPA task, mice were tested for memory retention in the CPA task. ChR2 mice that received HPC→AHN pathway stimulation during RTPA at 6 Hz, but not 20 Hz, displayed a robust conditioned aversion to the stimulation chamber (Fig. 5h). The distance travelled was not different between controls and ChR2 groups (Fig. 5i).

**Figure 5.**
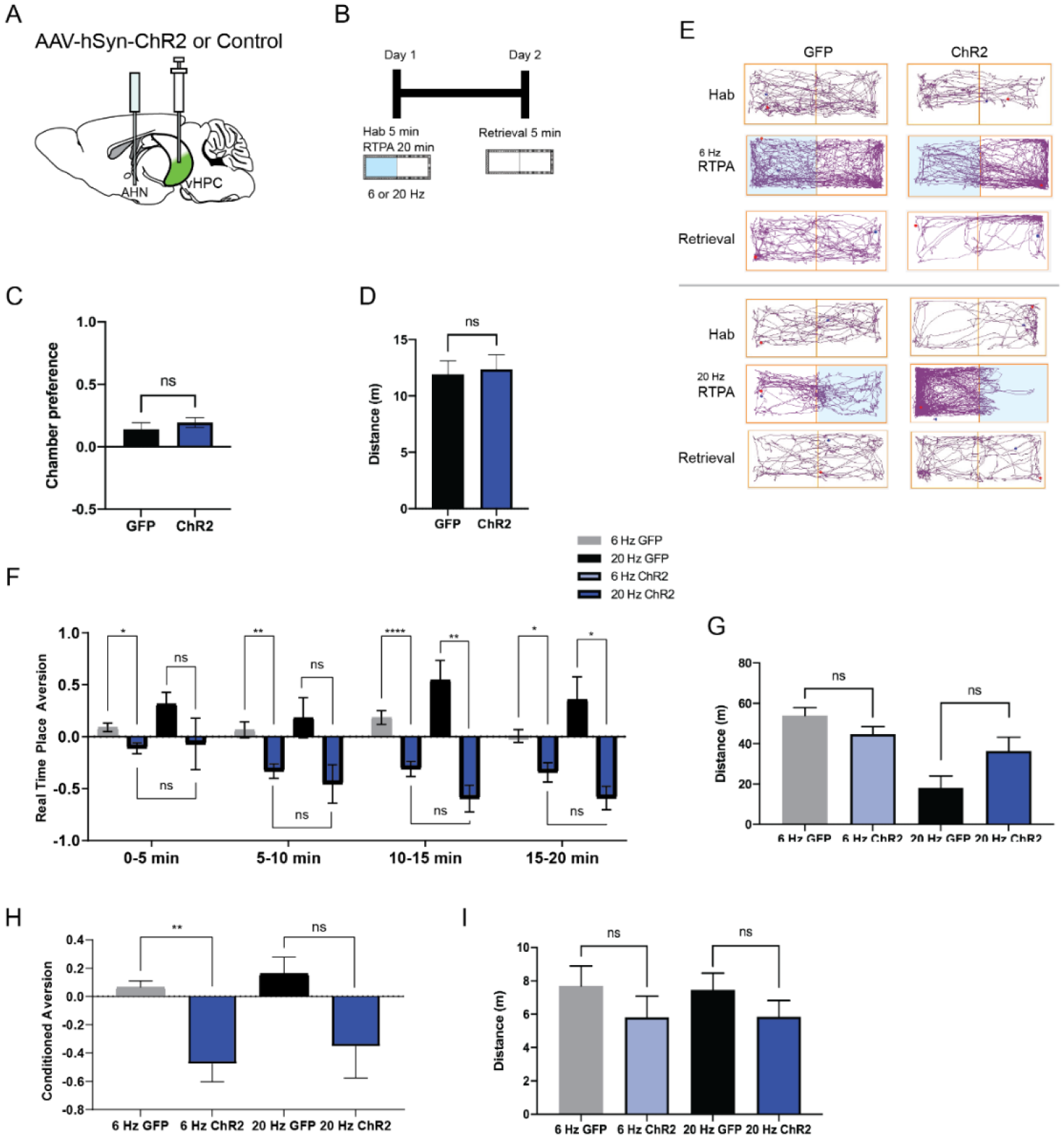
HPC→AHN pathway activation is aversive and instructs learning of a conditioned place aversion. **a,** Schematic illustration of optogenetic activation of hippocampal terminals in the AHN (GFP N= 21, ChR2 N=22). **b,** Schematic describing the RTPA and CPA test paradigm: day 1 consisting of habituation and real-time place preference (20 min) and day 2 for testing conditioned place preference (5 min). **c,** Chamber preference during habituation (unpaired t-test, two-tailed, t=0.8339, df=41, *p=0.4089*, NS). **d**, Distance travelled during habituation (unpaired t-test, two-tailed, t=0.2674, df=41, *p=0.7905*, NS). **e,** Representative locomotion trajectory for a GFP control animal (left column) and a ChR2-expressing animal (right column) during habituation (hab), 6 Hz or 20 Hz real-time stimulation (6 Hz RTPA, 20 Hz RTPA), and conditioned place aversion test (Retrieval). Light-coupled chambers are shown in blue. **f,** Real time place aversion monitored across 20 minute test. GFP 6 Hz vs. ChR2 6 Hz (2-WAY RM ANOVA, time x treatment F(3,120)=3.539, *\*p=0.0168*), GFP 20 Hz vs. ChR2 20 Hz (2-WAY RM ANOVA, time x treatment, F(3,30)=4.132, *\*p=0.0145*). GFP 6 Hz vs. GFP 20 Hz (2-WAY RM ANOVA, time x frequency, F (3, 75) = 1.249, *p=0.2980*, NS). ChR2 6 Hz vs. ChR2 20 Hz (2-WAY RM ANOVA, time x frequency, F (3, 75) = 1.828, *p=0.1492*, NS) **g,** Distance travelled during 6 Hz and 20 Hz real-time stimulation. GFP 6 Hz vs. ChR2 6 Hz (unpaired t-test, two-tailed, t=1.688, df=41, *p=0.0859*). GFP 20Hz vs. ChR2 20 Hz (unpaired t-test, two-tailed, t=1.507, df=10*, p=0.1627*, NS). **h,** Conditioned aversion memory tested 24-hours after real time place aversion tests. GFP 6 Hz vs. ChR2 6 Hz (unpaired t-test, two-tailed, t= 3.916, df=10, *\*\*p=0.0029*). GFP 20 Hz vs. ChR2 20 Hz (unpaired t-test, two-tailed, t=1.996, df=10, *p=0.0739*, NS). ChR2 6 Hz vs. ChR2 20 Hz (unpaired t-test, two-tailed, t=0.4771, df=10*, p=0.6436*, NS). GFP 6 Hz vs. GFP 20 Hz (unpaired t-test, two-tailed, t= 0.7483, df= 10, *p=0.4715*, NS). **i,** Distance travelled during the conditioned place aversion test. GFP 6 Hz vs. ChR2 6 Hz (unpaired t-test, two-tailed, t=1.095, df=10, *p=0.2991*). GFP 20 Hz vs. ChR2 20 Hz (unpaired t-test, two-tailed, t=1.153, df=10, *p=0.2756,* NS). All results reported are mean ± s.e.m. *p < 0.05, **p < 0.01, ***p < 0.001, ****p<0.0001.

### Optogenetic inhibition of HPC→AHN pathway impairs the retrieval of contextual memory of predator cue

The HPC→AHN pathway activity is aversive and can induce a conditioned place aversion. However, the nature of information that the pathway encodes remains unknown. Given the role of the HPC in contextual memory and its direct connection with the AHN, we hypothesized that the HPC→AHN pathway may promote goal-directed escapes by encoding the animal’s knowledge or memory of the surrounding environment.

To address this hypothesis, we first investigated the role of HPC→AHN pathway in mediating contextual memory to predatory threats by optogenetically inhibiting the pathway and measuring its effects on conditioned escape responses from an ethologically relevant predator cue. The HPC was virally transduced with AAV-CamKIIa-ArchT-GFP, and optic fibers were bilaterally implanted at the AHN to illuminate HPC axon terminals (Fig. 6a, Extended Data Fig. 1e,f). On day 1 (pre-conditioning), mice were habituated to two neutral but visually distinct contexts in a two-chamber apparatus. On day 2 and 3 (conditioning), a predator cue (10% L-Felinine) was paired with one chamber, and water with the other in a counterbalanced manner (Fig. 6b). L-Felinine is a putative predator kairomone of a Felidae species^44, 45^ and was chosen as a predator cue because it induces a robust dose-dependent increase in freezing compared to predator urine samples (Extended Data Fig. 6). During the conditioning, both GFP and ArchT mice displayed increased freezing in the L-Felinine chamber compared to the water chamber (Extended Data Fig. 7). On Day 4 (post-conditioning), mice were allowed to freely explore the two chambers while the HPC→AHN pathway was optogenetically inhibited (Fig. 6b). As expected, GFP control mice displayed an avoidance of the L-Felinine context (Fig. 6c,d, Extended Data Fig. 8a). ArchT mice, however, failed to remember and avoid L-Felinine context (Fig. 6c,d, Extended Data Fig. 8b). Furthermore, the contextual memory impairment was accompanied by significant decreases in defensive behavioural responses such as freezing, escape runs, and grooming compared to GFP control (Fig. 6e-h, Supplementary Movie 6). This finding was replicated in a different CPA paradigm involving 5 days of conditioning, which allowed us to quantify learning of predator context across multiple days (Extended Data Fig. 9a). Both GFP and ArchT mice developed predator odour context aversion gradually (Extended Data Fig. 9b,c), and the HPC→AHN pathway inhibition post conditioning resulted in contextual memory impairment (Extended Data Fig. 9d-f).

**Figure 6.**
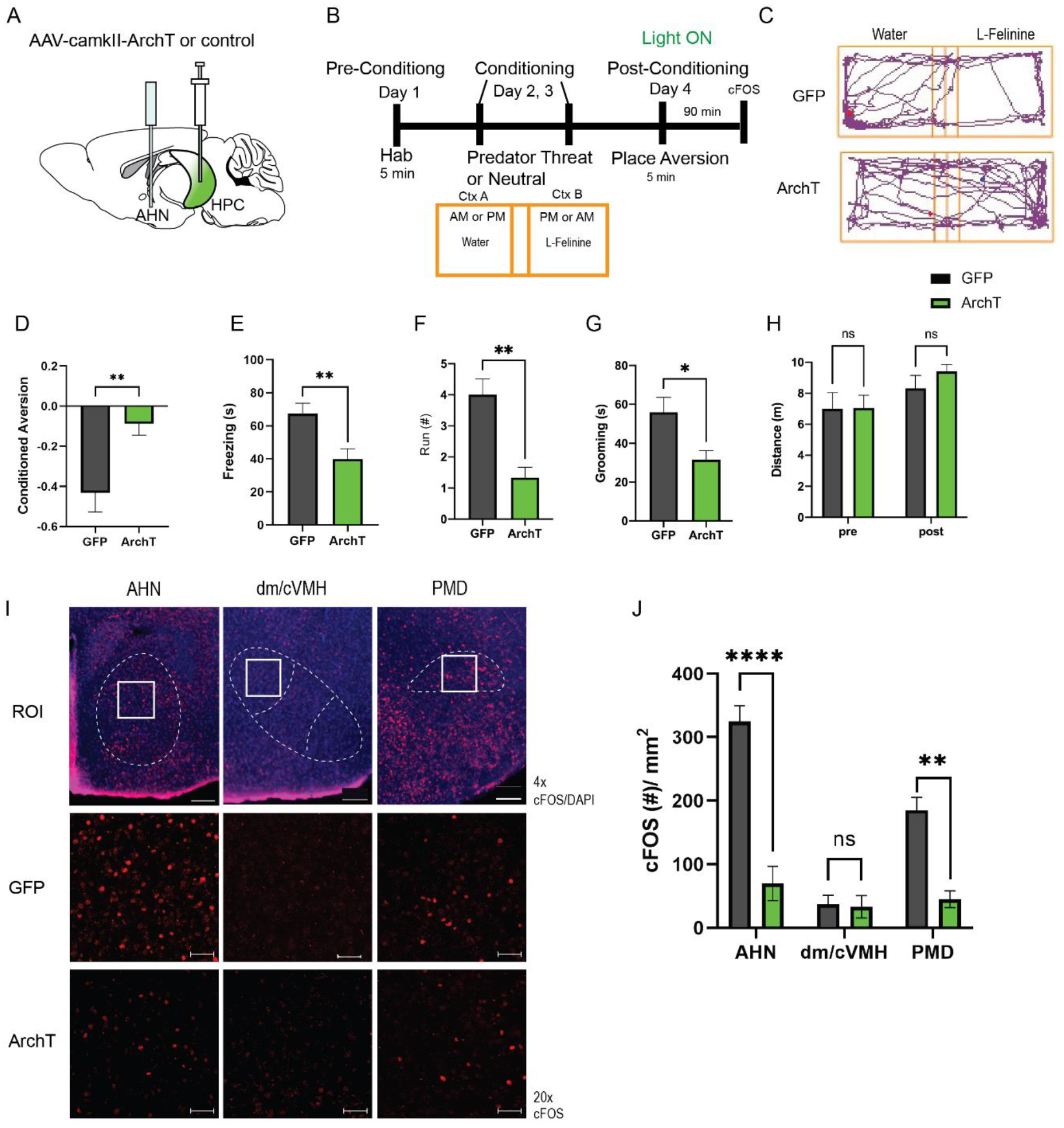
HPC input to the AHN is necessary for remembering the context-associated with predatory threat. **a**, Schematic illustration of optogenetic HPC terminal inhibition in the AHN (GFP N=8, ArchT N=12). **b**, Schematic describing the behavioural paradigm for the contextual fear conditioning with the predator odor (L-Felinine-); day 1 for habituation (5 min), days 2-3 for two daily conditioning sessions where mice were enclosed either L-Felinine- or water-paired chamber for 20 minutes in AM and PM in a counterbalanced manner, and day 4 for testing conditioned place preference (5 min) and the immunochemical detection of c-Fos. **c**, Representative locomotion trajectory for a GFP control animal (top) and a ChR2-expressing animal (down) during the conditioned place aversion test with optogenetic HPC terminal inhibition in the AHN (left: water-coupled chamber, right: L-Felinine-coupled chamber). **d**, Conditioned aversion memory tested 24-hours after conditioning. GFP vs. ArchT (unpaired t-test, t=3.223, df=18. *\*\*p=0.0043*). **e**, Freezing time during the conditioned place aversion test. GFP vs. ArchT (unpaired t-test, t=3.056, df= 18, *\*\*p=0.0062*). **f,** Number of escape runs from the L-Felinine-paired chamber to the water-paired chamber. GFP vs. ArchT (unpaired t-test, t=4.479, df=18, *\*\*\*p=0.0002*). **g,** Time spent grooming during the conditioned place aversion test. GFP vs. ArchT (unpaired t-test, t=2.816, df=18, *\*p=0.0107*). **h,** Distance travelled during habituation (pre) and conditioned place aversion test (post) (2-WAY ANOVA, training x treatment, F(1,20)=0.9938, *p=0.3307*, NS). **i,** c-Fos immunochemical detection across the medial hypothalamic defense system (AHN, VMHdm/c, PMD). First row, representative 4x epi-fluorescence microscope images of the medial hypothalamic defense system in GFP control mice. The regions of interest (ROI, white squares) within AHN, VMHdm/c, and PMD were imaged by confocal microscopy for cell counting. Second and third row: representative 20x confocal images of c-Fos signals in AHN, VMHdm/c, and PMD activated by the conditioned place aversion test in GFP and ArchT mice, respectively. **j**, Density of c-Fos signals in AHN, VMHdm/c, and PMD in GFP control (black) and ArchT mice (green) (N=5 mice for each group). GFP-AHN vs. ArchT-AHN (unpaired t-test, two-tailed, t=7.005, df=6, *\*\*\*p=0.0004*). GFP-VMHdm/c vs. ArchT VMHdm/c (unpaired t-test, two-tailed, t=0.5414, df=6, *p= 0.6078*, NS). GFP-PMD vs. ArchT-PMD (unpaired t-test, two-tailed t=5.810, df=6, *\*\*p=0.0011*). All results reported are mean ± s.e.m. *\*p < 0.05, **p < 0.01, ***p < 0.001, ****p<0.0001*. Scale bar = 100 μm for 4x epi-fluorescence microscope images) and 10 μm for 20x confocal images.

Next, we investigated the role of HPC inputs in driving the activities of AHN during the retrieval of contextual memory of predator cues. 90 min after a post-conditioning test, GFP control and ArchT mice were euthanized for immunochemical detection of c-Fos in the medial hypothalamic defense system. We found that c-Fos expression in ArchT mice was decreased in the AHN and PMD, but not in the VMHdm/c, compared to GFP control (Fig. 6i,j). Together, our data demonstrates that the HPC→AHN pathway enables animals to avoid the environment associated with predators by driving AHN activities during the retrieval of contextual memory of predator cues.

### Optogenetic activation of HPC→AHN pathway evokes goal-directed escapes to shelter

Another important aspect of escape response, other than predatory threats, is the use of shelter as the escape target. Thus, we tested whether the HPC→AHN pathway plays a role in goal-directed escape to a safe shelter. The HPC was virally transduced with AAV-hSyn-ChR2-GFP, and optic fibers were bilaterally implanted at the AHN (Fig. 7a). Mice were then placed in the open field arena containing a shelter box to determine whether the optogenetic pathway stimulation leads to an escape flight to the shelter (Fig. 7c). During the habituation stage, mice were given 5 minutes to freely explore the arena and exploit the shelter (Fig. 7b). Both ChR2 and GFP control mice intermittently visited the shelter and spent a comparable amount of time in the shelter (Fig. 7d,e). During the stimulation stage, the HPC→AHN pathway was stimulated at 6 or 20 Hz frequency when mice were outside the shelter. The pathway stimulation in ChR2 mice at both frequencies evoked goal-directed escapes toward the shelter, with a shorter latency to escape and a greater speed of escape running compared to GFP controls (Fig. 7f-i, Supplementary Movie 7,8). Thus, our findings show that the behavioural response evoked by HPC→AHN pathway activation is not just a simple increase in locomotion but constitutes a goal-directed escape toward a safe shelter.

**Figure 7.**
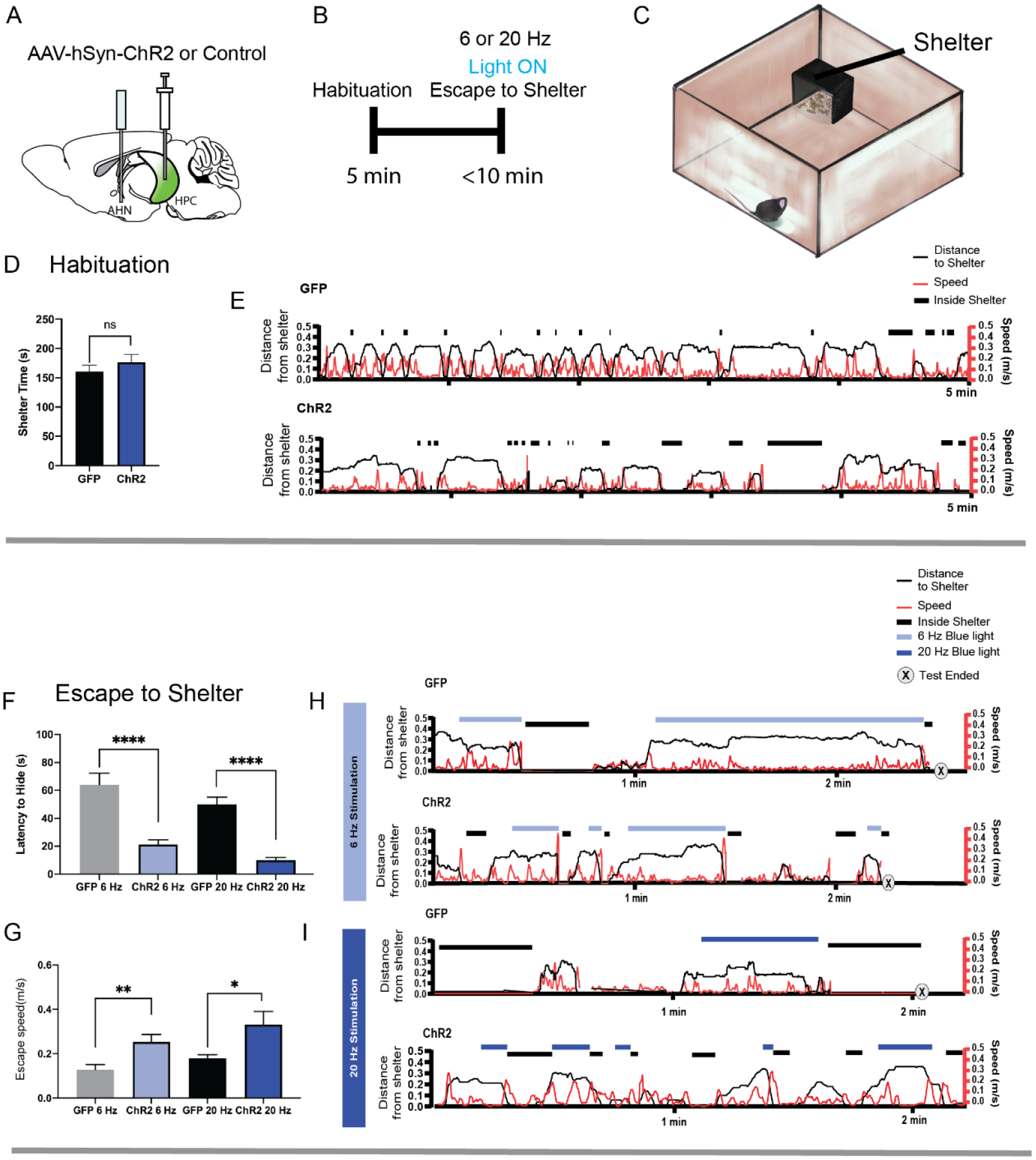
HPC→AHN pathway activation induces goal-directed escape. **a**, Schematic illustration of optogenetic HPC terminal activation in the AHN. **b**, Schematic describing a test paradigm consisting of habituation (5 min) and a 6 or 20 Hz stimulation stage to induce shelter-directed escapes. **c**, A cartoon drawing of the open field arena with a shelter box. **d**, Time spent in the shelter during habituation (GFP N=8, ChR2 N=6, unpaired t-test, two-tailed, t=0.9241, df=12, *p=0.3736*, NS). **e**, Representative line graphs for GFP (top) and ChR2 (bottom) mice, showing distance from shelter (black lines), speed (red lines), and moments when mice were inside the shelter (black boxes) over the 5 min habituation period. **f**, Latency to escape to the shelter after optogenetic HPC terminal activation. GFP 6 Hz vs. ChR2 6 Hz (GFP N=20, ChR2 N=19, unpaired t-test, two-tailed, t=4.631, df=36, *\*\*\*\*p<0.0001*). GFP 20 Hz vs. ChR2 20 Hz (GFP N=8, ChR2 N=10, unpaired t-test, two-tailed, t=7.733, df=16, *\*\*\*\*p<0.0001*). **g**, Speed of escape running. GFP 6 Hz vs.ChR2 6 Hz (GFP N=20, ChR2 N=19, unpaired t-test, two-tailed, t=3.092, df=37, *\*\*p=0.0038*). GFP 20 Hz vs. ChR2 20 Hz (GFP N=8, ChR2 N=11, unpaired t-test, two-tailed, t=2.167, df=18, *\*p=0.0439*). **h, i**, Representative line graphs for GFP and ChR2 mice, showing distance from shelter (black lines), speed (red lines), and moments when mice were inside the shelter (black boxes) during the 6 Hz (**h**) and 20 Hz (**i**) HPC terminal stimulation stage. Light and dark blue highlights indicate the duration of 6 Hz and 20 Hz light stimulation, respectively, and (x) denotes test termination time. All results reported are mean ± s.e.m. \**p < 0.05, **p < 0.01, ***p < 0.001, ****p<0.0001*.

### Optogenetic inhibition of HPC→AHN pathway impairs goal-directed escapes to shelter

Like predator odours, high frequency (17-22 kHz) ultrasound stimuli evoke strong defensive responses in mice, including escape flight and freezing. They are thought to serve as alarm cries to warn conspecifics of impending danger in response to a predator threat. For example, rats emit ultrasonic vocalization when approached by a cat, and the intensity of the alarm cries increases as a function of threat proximity ^46, 47^. A recent study found that the same ultrasound stimulus elicits different defensive responses depending on the availability of a safe shelter; mice display escape flights when a safe shelter is available but freezing when there is no shelter^6^. The study suggests that animals’ spatial knowledge about escape routes and shelter availability determines the best course of defensive actions in the face of predatory threats.

Given the role of animals’ spatial knowledge in shaping escape responses, we tested whether the HPC→AHN pathway activity is necessary for mice to use mnemonic information about shelter availability and location during the ultrasound-evoked escape (Fig. 8a,b). During a habituation stage (7 min), mice were allowed to explore a modified Barnes maze with 20 equally spaced holes, one of which leads to a shelter box (Fig. 8b,c). Both ArchT and GFP control mice found the shelter at least once during the survey stage and spent a comparable amount of time in the shelter (Fig. 8d), indicating that the two groups had a similar condition to memorize shelter location and shelter availability. During the subsequent threat delivery stage, light illumination (i.e., pathway inhibition) started right after mice voluntarily came out of the shelter, and a 9 s ultrasound stimulus (20 kHz) was triggered manually at randomized positions on the platform. Upon hearing the ultrasound threat, mice first turned its heads towards the shelter and initiated an escape flight, reaching a maximum running speed at the middle of the escape path, which are the known features of the goal-directed escape flight (Fig. 8e, shelter, Extended Data Fig.10 a-c). Consistent with a previous report, mice displayed these characteristic escape responses only when a shelter is available. When the shelter was removed from the maze before the survey stage, the ultrasound threat failed to elicit escape flights, but instead caused either freezing or slow and disorganized flights in random directions (Fig. 8e, no shelter, Extended Data Fig. 10d-h).

**Figure 8.**
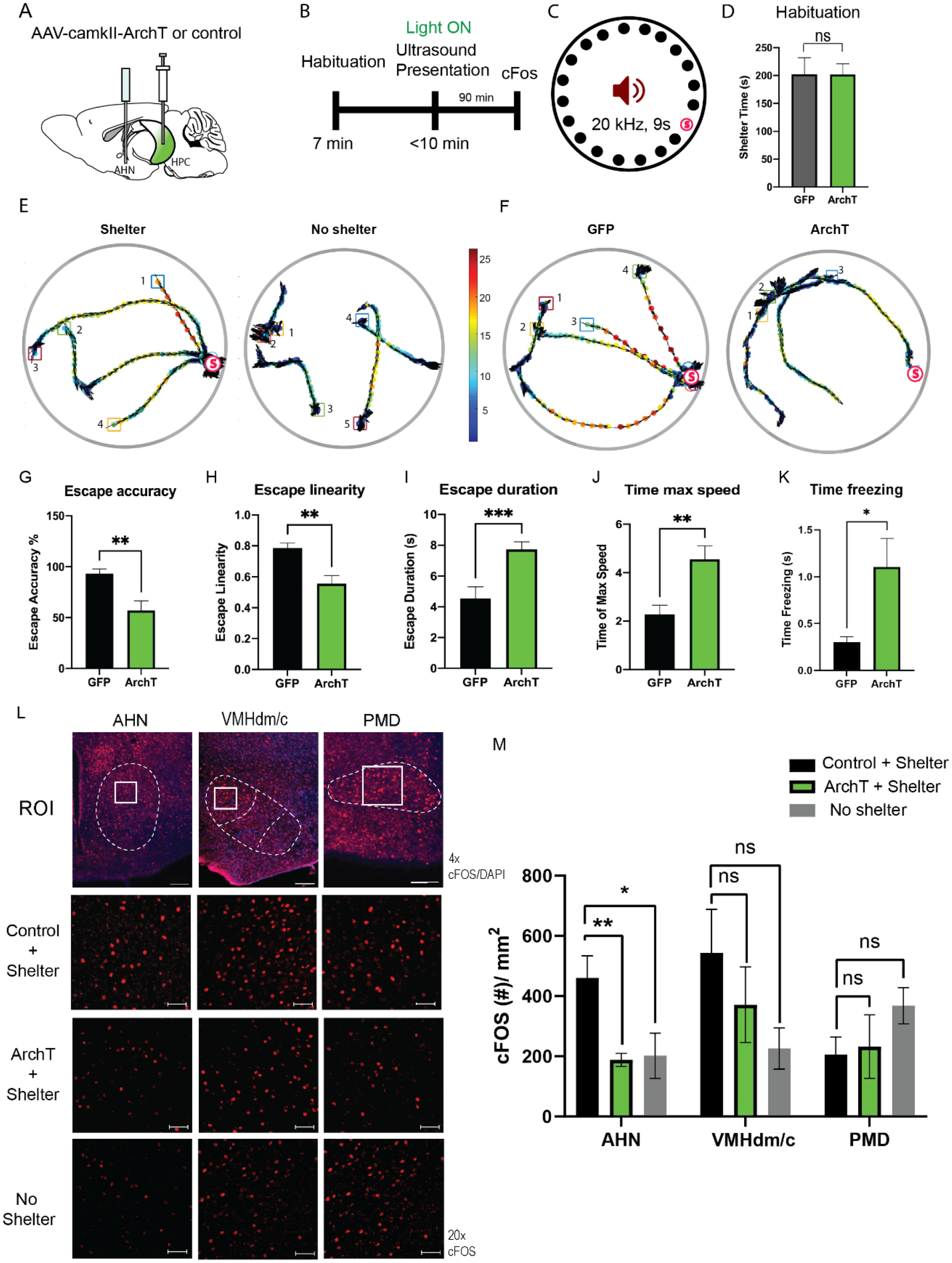
HPC→AHN pathway is necessary for goal-directed escape. **a,** Schematic illustration of optogenetic HPC terminal inhibition in the AHN (GFP N=7 and ArchT N=15). **b**, Schematic describing a test paradigm consisting of habituation (7 min) and a threat delivery stage during which ultrasound (20 kHz, 9s) is turned on after mice voluntarily come out of the shelter to induce a shelter-directed escape. **c**, Top view of testing apparatus, a modified Barnes maze. “s” denotes the position where a shelter was placed. The red speaker sign denotes the auditory threat played from a speaker above the apparatus centre. **d**, Time spent in the shelter during habituation stage (unpaired t-test, t=0.01343, df=21, p=0.9894, NS). **e**, Representative ultrasound-evoked escape trajectories for a Wild type (WT) when a shelter is available (left) vs. WT when no shelter is available (right). **f**, Representative ultrasound-evoked escape trajectories for a GFP control (left) and ArchT (right). **e,f,** Individual threat presentation as a trial is numbered next to a square box, which denotes the animals’ starting position at the beginning of 9 s of 20 kHz sound. Dot color along the trajectory lines reflects animals’ speed. The arrows track animals’ head direction. The heatmap colorbar displays the scale of speed (pix/s). **g**, Accuracy of reaching the shelter during escape. (unpaired t-test, t=3.149, df=33, *\*\*p=0.0034*). **h**, The linearity of escape trajectories expressed as the percentage ratio between the length of escape trajectory and a linear distance from escape onset position (i.e., ultrasound onset) to the shelter (unpaired t-test, t=3.266, df=31, *\*\*p=0.0027*). **i**, Time elapsed from the ultrasound onset to the shelter arrival (unpaired t-test, t=3.666, df=33, *\*\*\*p=0.0008*). **j**, Time elapsed to reach the maximum speed during escape running to the shelter (unpaired t-test, t=3.134, df=31, *\*\*p=0.0036*). **k**, Time spent in freezing between the ultrasound onset and the shelter arrival (unpaired t-test, t=2.261, df=33, *\*p=0.0305*). **l**, c-Fos immunochemical detection across the medial hypothalamic defense system (AHN, VMHdm/c, PMD). First row, representative 4x epi-fluorescence microscope images of the medial hypothalamic defense system in GFP control mice. The regions of interest (ROI, white squares) within AHN, VMHdm/c, and PMD were imaged by confocal microscopy for cell counting. Second and third row: representative 20x confocal images of c-Fos signals in AHN, VMHdm/c, and PMD activated by ultrasound-evoked escapes with a shelter available in GFP (second row) and ArchT mice (third row). Fourth row: representative 20x confocal images of c-Fos signals activated by ultrasound-evoked escapes without shelter in controls. **m**, Density of c-Fos signals activated by ultrasound-evoked escapes in AHN, VMHdm/c, and PMD in GFP controls with shelter (black), ArchT mice with shelter (green), and controls without shelter (grey)(N=5 mice for each group). GFP-AHN vs. ArchT-AHN (unpaired t-test, two-tailed, t=7.005, df=6, *\*\*\*p=0.0004*). GFP-VMHdm/c vs. ArchT VMHdm/c (unpaired t-test, two-tailed, t=0.5414, df=6, *p= 0.6078*, NS). GFP-PMD vs. ArchT-PMD (unpaired t-test, two-tailed t=5.810, df=6, *\*\*p=0.0011*). AHN: control + shelter vs. ArchT + shelter (unpaired t-test, two-tailed, t=3.567, df=8, *\*\*p=0.0073*). AHN: control + shelter vs. control with no shelter (unpaired t-test, two-tailed, t=2.467, df=8, *\*p=0.0389*). VMHdm/c: control + shelter vs. ArchT + shelter (unpaired t-test, two-tailed, t=0.8972, df=8*, p=0.3958*, NS). VMHdm/c: control + shelter vs. control with no shelter (unpaired t-test, two-tailed, t=1.98, df=8, p=0.0830, NS). PMD: control + shelter vs. ArchT + shelter (unpaired t-test, two-tailed, t=0.2159, df=8, *p=0.8345*, NS). PMD: control + shelter vs. control with no shelter (unpaired t-test, two-tailed, t=1.937, df=8, *p=0.088*, NS). Scale bar = 100 μm for 4x epi-fluorescence microscope images) and 10 μm for 20x confocal images.

Optogenetic inhibition of the HPC→AHN pathway produced a range of effects on the goal-directed escape. Instead of a quick and direct flight to shelter seen in GFP controls, ArchT mice displayed disorganized escape trajectories and slow escape running speed, reminiscent of how the control mice respond to ultrasound when the shelter is not available. (Fig 8f, Extended Data Fig.11 a,b, Supplementary Movie 9). In addition, ArchT mice directed their flights to locations farther away from the target shelter (i.e., lower escape accuracy, Fig. 8g), resulting in low escape success rate of 30 % (6 out of 20 trials) compared to 87% in GFP controls (13 out of 15 trials) (Extended Data Fig. 11 c,d). The lower escape accuracy and rate of successful escape suggest that ArchT mice failed to use a memory of shelter availability and shelter location to support their goal-directed escapes. Furthermore, a decrease in the organization and efficiency of escape was indicated by changes in various parameters such as escape linearity, escape duration, time to reach the maximum speed and increased freezing (Fig. 8h-k). The impairments in goal-directed escape were not accompanied with any changes in anxiety-related behaviours (Extended Data Fig. 12).

Lastly, we investigated the role of HPC inputs in driving AHN activities during the goal-directed escape. ArchT mice were exposed to an ultrasound-evoked escape flight coupled with the HPC→AHN pathway inhibition (ArchT/Shelter+). Control mice were split into two groups, where one group was exposed to an ultrasound-evoked escape flight with a shelter available (Control/Shelter+) and the other group without shelter (Control/ Shelter-). 90 min later, mice were euthanized for immunochemical detection of c-Fos. We found that when the HPC→AHN pathway was inhibited (ArchT/Shelter+), c-Fos expression was significantly reduced in the AHN, but not in the VMHdm/c and PMD, compared to controls (Control/Shelter+) (Fig. 8l,m). Furthermore, upon the removal of shelter, c-Fos expression in the AHN of control mice decreased to the level of ArchT mice (Fig. 8l,m), suggesting that the AHN is not activated properly during the ultrasound-evoked escape if the HPC→AHN pathway is inhibited, or if a shelter is not available. Thus, the inhibition of HPC→AHN pathway and the removal of shelter during the ultrasound-evoked escape produced similar effects not only at the behavioural level but also on the AHN activity. Taken together, these results demonstrate that the HPC→AHN pathway supports the goal-directed escape by driving the AHN activity with mnemonic information about shelter availability and shelter location.

## Discussion

Current knowledge of HPC control of fear and defensive response has largely been derived from studies of associative memory for nociceptive stimuli (e.g., electric foot shocks)^26, 48, 49^. While informative, they have left a widening gap between our understanding of the neural circuit mechanism underlying fear and the complex innate defensive behaviours displayed in natural environments. Our investigation of the HPC-AHN pathway provides a framework on how explicit memory and transmission of contextual information control innate defensive behaviours. To our knowledge, the present study is the first to 1) delineate a direct functional connection between the hippocampus and the medial hypothalamic defense system and to 2) show how hippocampal signals representing environmental contexts control innate defensive responses at the neural circuit level.

We found that a direct optogenetic stimulation of the entire AHN structure increases avoidance, immobility, and escape running and jumping. This is consistent with a recent finding that the selective activation of VMHdm/c inputs to the AHN elicits escape running, immobility, and jumping^12^. Interestingly, however, the selective activation of VMHdm/c inputs to the dorsolateral periaqueductal gray produced only immobility but failed to evoke escape running and jumping. These findings, together with ours, support the idea that distinct aspects of defensive responses to threat are mediated by different cell types and efferent projections in the VMHdm/c and AHN. The AHN has been shown to be a largely GABAergic structure, with some scattered glutamatergic cells located at the ventral aspect of the medial zone^50, 51^. Thus, it remains to be investigated whether VMHdm/c inputs selectively target AHN GABA or glutamatergic cells, or both. Of note, we did not observe any escape jumping upon stimulating AHN GABA cells (data not shown). We speculate that the activity of AHN glutamatergic, but not GABAergic, cells may be sufficient to evoke escape jumping, and that the escape jumping induced by the VMHdm/c-AHN pathway may have been driven at least in part by inputs to AHN glutamatergic cells.

Several tract-tracing studies, including ours, have shown that the medial hypothalamic defense system receives strong excitatory inputs from the hippocampus^36, 52^. Our anterograde tracing experiments revealed that hippocampal axon terminals are found almost exclusively at the AHN, but not the PMD and VMHdm/c. This suggests that the AHN is the main entry site for hippocampal signals in the hypothalamic defense system, therefore an ideal brain region for integrating environmental context and concomitant predator sensory information to support the contextual memory of predator threats. In line with this idea, a recent work found that predatory context (e.g., cat-associated context) induces a robust increase in c-Fos level in the AHN^53^. Interestingly, the study identified the PMD, not the AHN, as the most responsive hypothalamic region to predatory context, despite the relative scarcity of hippocampal innervation in the PMD. It is possible that the AHN and the PMD act together to support the contextual memory of predator threat, where the AHN first receives contextual information from the hippocampus and then conveys it to the PMD. This possibility will have to be tested by selectively blocking AHN inputs to the PMD and analyzing its impact on the contextual memory of predatory threats.

Electrophysiological recordings confirmed the monosynaptic nature of the HPC→AHN connection. In slice recordings experiments, patching was guided by mCherry fluorescence in double transgenic (Dlx5/6-Flpe; Frepe) reporter mice. The Dlx5/6-Flpe line has been used to label GABA cells in the forebrain cortex and hippocampus with a high labelling efficacy and specificity^54–58^. Optogenetic stimulation of hippocampal axon fibers in the AHN evoked robust EPSPs in both GABAergic and non-GABAergic cells with an onset latency less than 5 ms that indicates monosynaptic responses. It should be noted, however, the labelling efficacy of the Dlx5/6-Flpe mouse line has not been thoroughly characterized in the hypothalamic regions, including the AHN. Thus, our study may overestimate the abundance of non-GABAergic AHN cells receiving monosynaptic hippocampal inputs. Experiments using other reporter strains such as GAD67-GFP mice will help to further clarify the abundance of hippocampal inputs to non-GABAergic cells in the AHN.

Compared to a direct AHN stimulation which invariably induced robust escape jumping, HPC→AHN pathway activation only increased running bouts and speed. This suggests that escape jumping likely requires additional inputs to the AHN from other structures such as amygdala and VMH which carry sensory information about predatory threats. On the other hand, however, the activation of HPC→AHN pathways was as powerful as direct AHN stimulation in producing a strong real-time and conditioned place aversion, suggesting that the pathway can form a lasting memory of threat-associated environmental context. Indeed, we found that upon the inhibition of HPC→AHN pathway, mice failed to remember where a predator cue was previously encountered. This indicates that the hippocampus controls the medial hypothalamic defense system and mediates the contextual memory of predator threats via its direct projections to the AHN. Aversive experience such as the predator odour can induce remapping of place cell firing which becomes stabilized after 24 hours. Thus, HPC→AHN pathway is likely activated by a specific hippocampal place cell ensemble that represents predator odour location and context. It is of note, however, that our optogenetic inhibition targeted only the retrieval phase of memory in the conditioned place aversion. Thus, it remains unknown if the HPC→AHN pathway is also involved in the memory encoding and how the synaptic plasticity of the HPC inputs to AHN changes during memory encoding and reconsolidation.

HPC→AHN pathway activation evoked goal-directed escapes, whereas its inhibition disrupted a successful escape to shelter. In the ultrasound-evoked escape assay, mice detect an ultrasound threat and then evaluate whether shelter is available based on their memory of shelter. If shelter is available, mice compute the flight direction before launching an escape, and if not, they freeze. Importantly, a previous study using the same assay has shown that unlike in Morris water maze or Barnes maze tests, mice can make accurate escape flights even in complete darkness, suggesting that external landmarks (reference memory) are not required for mice to determine the shelter location. Instead, mice compute an escape vector to the shelter location by integrating self-motion over time using the path integration strategy^6, 59, 60^. We found that instead of making a quick and direct escape to shelter, ArchT mice display a slow escape running and are more likely to flee to unsafe locations away from the target shelter. The observed behavioural impairments present multiple possibilities regarding the role of HPC→AHN pathway. The most plausible explanation is that the pathway uses mnemonic information about the shelter availability and shelter location to increase a motivational drive to escape. When mice hear the ultrasound threat, the hippocampus may reactivate shelter memory and send the signals to the AHN, thereby increasing the escape drive and escape-associated locomotion. If the pathway is optogenetically inhibited, however, the shelter memory recall would no longer be able to activate the AHN and support goal-directed locomotion, resulting in a slow or even lack of escape running. Alternatively, the HPC→AHN pathway may represent not only a motivational drive to escape but also encode specific geometric information about a shelter location generated by hippocampal place cells. Such information may be used for the path integration process either within the medial hypothalamic defense system or its downstream targets to compute an escape vector.

## Methods

### Animals

Adult C57BL/6 mice (Charles river) at 8-12 weeks of age were used for AHN soma activation and HPC terminal activation studies. Double transgenic Dlx5/6-FLPe;RC::FrePe mice by crossing homozygous RC::FrePe^52^ mice were crossed with Dlx5/6-FLPe mice [Tg(mI56i-FLPe)39Fsh/J, JAX#010815]. 10 days prior to testing, animals were single housed with food and water provided *ad libitum* in 12 hr light/dark cycle. All procedures were approved by the Local Animal Care Committee (LACC) at University of Toronto.

### Viral vectors and Stereotaxic surgery

AAV2/8-hsyn-hChR2 (H143R)-GFP, AAV2/5-camk2a-eArchT3.0-GFP, AAV2/8-hsyn-GFP, and AAV2/5-camk2a-GFP were purchased from the Addgene Viral Vector Core and used as received. For all surgical procedures, mice were anaesthetized with isoflurane (4% for induction and 2% for maintenance of anaesthesia) at an oxygen flow rate of 1 L/min, and head fixed in a stereotactic frame (David Kopf). Eyes were lubricated with an ophthalmic ointment throughout the surgeries.

Ketoprofen was provided for pain management during post-operative recovery. Viruses were infused by pressure injection. For the AHN infusion (AP −0.85mm, ML 0.45 mm, DV −5.2 mm), 69 nL per site was infused by a pulled glass needle and Nanojet II (Drummond Scientific) at 46 nl/s rate, and the needle was left in place for additional 10 minute to limit the virus drag during needle retract. For the ventral hippocampus/subiculum infusion (AP −3.8mm, ML −2.1mm, DV −4.8mm, 10° away from the midline), 300 nL per site were infused by cannula needle connected to Tygon tubing to a 10-µL Hamilton syringe (Hamilton Company) at rate 0.1 μl/ min. Custom made ferrule fibers consisting of optic fibers (200 µm core diameter, 0.39 NA, Thorlabs) threaded in 1.25 mm wide zirconia ferrules (Thorlabs) were implanted at the AHN (AP −0.85 mm, ML 1.38 mm, DV −5.1 mm, 10° towards the midline) 2 weeks after the viral infusion surgery. All animals were handled for a minimum of 5 minutes for 3 days prior to behavioural testing 2 weeks post implant operation.

### Optogenetic manipulation

For bilateral light delivery, a patch cable (200 μm core diameter, 0.37 NA; Doric Lenses) was connected to a 1 x 2 optical commutator (Doric Lenses) to divide the light path into two arena patch cables attached to the implanted optic fibers. For ChR2-mediated optogenetic stimulation, blue light (473 nm, 6 Hz or 20 Hz) was produced using an arbitrary waveform generator (Agilent, 33220A) and a diode-pumped solid-state laser (Laserglow) at a power intensity of 5 mW from the optic fiber tip. For ArchT-mediated optogenetic inhibition, green light (532 nm, Laserglow) was applied continuously at a power intensity of 15 mW from the optic fiber tip. Light power was measured at the optic fibre tip using a power meter (PM121D, Thorlab) before each behavioural test.

### Optostimulation-evoked escape responses in open field and physical restraint

Following a 5-min habituation to the tethering cable in home cage, animals were placed in a clear plexiglass chamber (short walled-escapable condition: 50cm x 50 cm x 20cm) or in an opaque walled chamber (inescapable condition: 30cm x 30 cmx 30cm). In the clear chamber, low and high frequency photostimulation effects were compared while keeping consistent light power (5 mW) at the optic fibre tip. Animals were given two 2-minute photostimulation (6 Hz, 5ms pulse width) each followed by two minutes off period to observe the light offset effect. Rearing, jumping, freezing, and grooming were manually scored by key press in ANY-MAZE (Stoelting Co) for all animals. 2 weeks after testing the effects of AHN stimulation in the chambers (condition easy and hard), animals were tested in a physical restraint. DecapiCones (Braintree Scientific) were cut around head and shaved neck to create spaces for arena cable linked to animals’ head cap and for collar sensor with a pulse oximeter (STARR Life Sciences). Animals were restrained in the DecapiCones during physical restraint and received AHN stimulation (473nm blue light, 20Hz, 10s ON, 10 s OFF) for 30 minutes.

### Real Time Place Aversion (RTPA) and Conditioned place Aversion (CPA)

A custom-made 45 cm × 20 cm × 35 cm apparatus was equally divided such that each side possessed a distinct visual context. After 5 minutes of habituation, the preferred chamber was selected as the stimulation chamber. Animals received either 6 or 20 Hz blue light illumination upon entering the stimulation chamber during a 20 minute RTPA test. 24 hours after the RTPA test, animals were re-introduced to the two-chamber apparatus with light off, and the preference during the first 5 minute was analysed to measure the retrieval of CPA memory. Animals were placed back into the home cage for 5 minute and reintroduced to the testing apparatus to begin a second RTPA and CPA with 6 or 20 Hz stimulation. The order of stimulation frequency was pseudorandomized. ANY-MAZE software was used to determine the amount of time spent in each chamber and their corresponding track plots. Conditioned place aversion assessed the change in place preference index before and after the conditioning session. Place preference indexes for pre and post conditioning were calculated as the normalized difference out of total chamber time when animal was in the context.

### Predator odour contextual fear conditioning

A custom-made 45 cm × 20 cm × 35 cm apparatus was equally divided such that each side possessed a distinct visual context. Two different testing paradigms were employed. Both tests consisted of “pre-conditioning” phase (5 min), a habituation period to the two-chamber apparatus. The preferred chamber was always paired with L-Felinine. All behavioral and tracking analysis were done using ANY-MAZE software. In the AM/PM conditioning design, mice were enclosed in either L-Felinine (10 %)- or water-paired chamber for odour-context pairing for 20 minutes in AM and PM in a counterbalanced manner for two days^61, 62^. On day 4, animals were placed back in the two-chamber apparatus and measured for defensive behaviors and chamber preference. The second conditioning paradigm (Extended Data Fig. 9) measured place aversion development after each predator odour-context pairing. A total of 6 predator odour-context pairings were carried out from day 0. Each day consisted of 5 minutes of free exploration, followed by a 6-minute L-Felinine (0.3%) pairing to the context of preferred chamber side. Aversion index was calculated as: Aversion index = [(Time spent in preferred side - Time spent in less preferred side)/Total chamber time]. Conditioned place aversion assessed the change in place aversion index before and after the L-Felinine conditioning sessions as: Conditioned place aversion index = Aversion index of Post-Conditioning – Aversion index of Pre-Conditioning. In AM/PM design, escape running was defined as an event in which mice left the L-Felinine-paired chamber with a peak running speed greater than 50% of the average ambulation speed during the test. Freezing and grooming behaviours were manually scored.

### Optostimulation-evoked goal-directed escape

Mice were introduced to a chamber (40 cm x 40 cm x40 cm) under a dim red light condition (10 lux). A shelter box (12 cm x 12 cm x 8 cm) was placed at a corner of the chamber with home cage bedding material placed inside the shelter box as an olfactory cue. After 5 minutes of habituation session, a 6 or 20 Hz photostimulation was delivered when the mouse body centre was a minimum 25 cm away from the shelter and the head was not pointing towards the shelter. Latency to escape was measured as the time (sec) elapsed from the light onset until mouse directed its head and started to move toward the shelter. Speed of escape was measured as the peak speed during escape flight.

### Ultrasound-evoked escape assay

The ultrasound-evoked escape assay, modified from Vale et al. (2017), was conducted under a dim red light condition (10 lux). The behavioural apparatus was a Barnes maze - a white plastic circular platform (92 cm in diameter) with 20 equally spaced holes (5 cm in diameter and 5 cm away from the border of platform) that are blocked by plastic covers. A plastic shelter box (9 cm x 12 cm x 9 cm) was placed at one of the 20 holes with home cage bedding material inside to serve as an olfactory cue. Animals were given a minimum 7 min for the habituation stage, but if they did not find the shelter, they were given an additional 5 min. The ultrasound stimulus (20 kHz sine waveform, 9s duration, 75 dB) was generated by an amplifier (Topaz AM10) and an ultrasound speaker (L60, Pettersson) positioned 50 cm above the arena. Overhead videos were obtained using a webcam and analysed using the DeepLabCut to track animals’ body parts (nose, centre, and tail base). Locomotion speed, head direction angle (0 ∼180 degree), and distance between shelter and body parts were calculated with custom-written Matlab scripts. A successful arrival at the shelter was counted when animal’s body centre was inside the shelter. Escape accuracy was calculated from how much the shelter target was missed (i.e., how far the animal body was away from the shelter at the end of the 9 second ultrasound stimulus) using an equation [Accuracy = 100% - 10% * (distance between body centre and shelter/shelter diameter)]. Freezing behaviours were manually scored.

### Electrophysiology

Brains were rapidly removed after decapitation and placed into a cutting solution containing the following (in mM): 87 NaCl, 2.5 KCl, 25 NaHCO_3_, 0.5 CaCl_2_, 7 MgCl_2_, 1.25 NaH_2_ PO_4_, 25 glucose and 75 sucrose (Osmolarity: 315–320 mOsm), saturated with 95% O_2_/5% CO_2_. Coronal sections (250 μm thick) containing the hypothalamus were cut using a vibratome (VT-1200, Leica Biosystems). The aCSF solution consisted of the following (in mM): 123 NaCl, 2.5 KCl, 1.25 NaH_2_PO_4_, 26 NaHCO_3_, 10 glucose, 2.5 CaCl_2_ and 1.5 MgCl_2_, saturated with 95% O_2_/5% CO_2_, pH 7.4, osmolarity 300 mOsm. Slices were recovered at 34 °C in artificial cerebrospinal fluid (aCSF) for 30 min and subsequently kept at room temperature. During experimentation slices were perfused at a rate of 2 ml/min in aCSF and maintained at 27-30 °C.

Borosilicate glass micropipettes (BF120-69-15, Sutter Instruments) were pulled in a Flaming/Brown Micropipette Puller (P-1000, Sutter Instruments) and filled with an intracellular fluid containing the following (in mM): 108 K-gluconate, 2 MgCl_2_, 8 Na-gluconate, 1 K_2_-ethylene glycol-bis(β-aminoethyl ether)-N,N,N^′^, N^′^ -tetraacetic acid (EGTA), 4 K_2_-ATP, 0.3 Na_3_-GTP, 10 HEPES (osmolarity: 283-289 mOsm and pH: 7.2-7.4). The resistance of the pipettes was between 3–5 MΩ. Inhibitory post-synaptic currents (IPSCs), and excitatory post-synaptic currents (EPSCs) were blocked using bath application of 100 μM picrotoxin and 10 μM DNQX, respectively. Action-potential-dependent synaptic activity was blocked using 1 μM TTX and monosynaptic release was recovered by subsequent application of 100 μM 4-AP. All recordings were performed on minimum of five animals per group. EPSCs were recorded in voltage-clamp mode with the membrane voltage held at −70mV. For cell-attached recordings, light stimulation was performed in 5 ms pulses of 473 nm blue light at 15Hz. Light evoked excitatory post-synaptic potentials (eEPSCs) were obtained with a 5 ms pulses of 473 nm light with inter stimulus interval at a rate of 2 Hz.

Whole cell patch clamp recordings were obtained using a Multiclamp 700B amplifier (Molecular Devices, California, USA), low pas filtered at 1 kHz and digitized at a sampling rate of 20 kHz using Digidata 1440A (Molecular Devices). Data was recorded on a PC using pClamp 10.6 (Molecular Devices) and analysed using Clampfit (Molecular Devices).

### Histology

For c-Fos immunohistochemistry, mice were anesthetized with avertin 90 minutes after an exposure to predator context retrieval or ultrasound-evoked escape and underwent transcardial perfusion with 0.1M phosphate-buffered saline (PBS, pH 7.4), followed by 4% paraformaldehyde (PFA). The brain tissues were removed and were immersed in 4% PFA overnight and cryoprotected in a 30% sucrose solution for 48 hrs. Free-floating coronal sections (40 μm) were cut with a cryostat (Leica, Germany), permeabilized with PBS containing 0.3% Triton X-100 (PBS-T) and blocked with 5% normal donkey serum (Jackson ImmunoResearch). The tissues sections were then incubated with primary antibody (rabbit anti-c-Fos, SC-52; Santa Cruz Biotechnology, 1:1000 in PBS-T) at 4°C for 72 hrs. Next, the sections were rinsed with PBS-T and incubated with PBS-T containing Alexa Fluor 594-conjugated donkey anti-rabbit secondary antibody (1:500, Jackson ImmunoResearch Laboratories) at room temperature for 2 hrs. Sections were then rinsed with PBS, mounted on slides, and stained with DAPI solution (1 µg/ml in PBS) before coverslipping. Confocal microscope z-stack images were captured using a 20x objective lens on a LSM800 microscope (Zeiss, Germany). The image processing and cell counting were performed using the ZEN 2.6 blue software. For the confirmation of AAV infusion and fiber implantation sites, brain tissues were sectioned as described above and stained for GFP (chicken anti-GFP, 1:1000 in 0.1% PBS-T, Abcam, ab 13970; Alexa Fluor 488-conjugated donkey anti-chicken secondary antibody, 1:1000 in 0.1% PBS-T).

### Statistical analysis

All statistical analysis was performed using GraphPad Prism (GraphPad Software). In behavioural experiments, a (two-tailed) unpaired Student’s t-test were generally used, but two-way repeated-measures ANOVA (2-WAY RM ANOVA) was employed in the RTPA analysis with treatment groups (GFP vs. ChR2) and stimulation frequency (6 and 20 Hz) as a between-subjects factor and time as a within-subjects factor. For HPC terminals quantification, one-way ANOVA was used with the *post-hoc* analysis of Dunett multiple comparison test. For secondary predator odour context conditioning, one-way repeated ANOVA was used and followed by Dunnett’s multiple comparison test. Where appropriate, 2-WAY RM ANOVAs were followed by planned pairwise comparisons such as Sidak’s multiple comparison. A simple linear regression analysis was used to detect the relationship between the dose-dependent change in investigation time vs. freezing. A non-linear fit was used to model the change in speed and head angle vs. the normalized distance from the shelter in US evoked shelter directed escape. Significance was defined as *p < 0.05, \*\**p < 0.01, ***p < 0.001, ****p* < 0.0001.

---

**Extended Data Fig 1.**
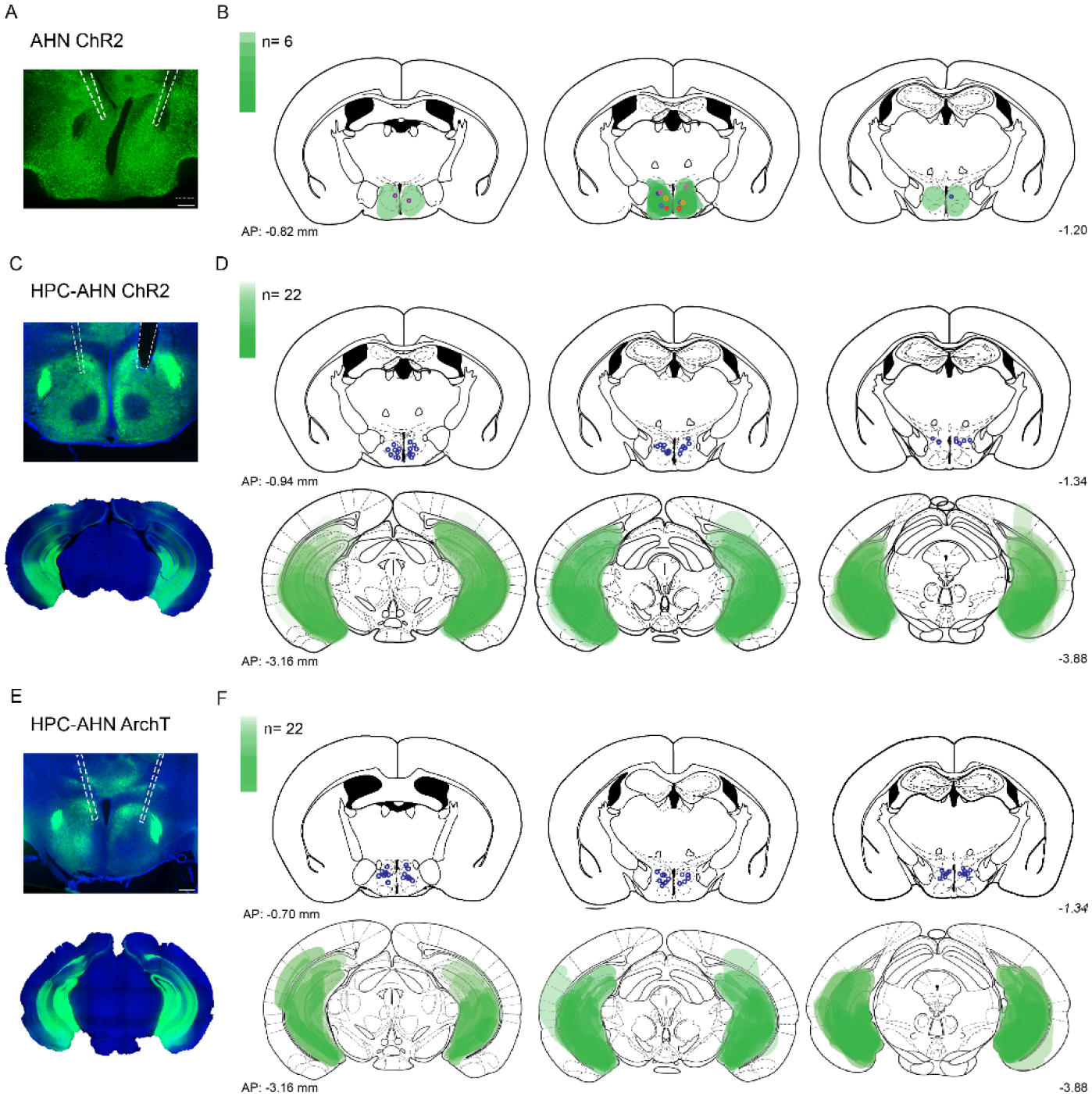
Compilation of viral expression and optic fiber implantation sites for optogenetic manipulation experiments. **a,b,** AHN soma activation. (N=6, each pair of colored circles shows the optic fiber tip placements in one animal). **c,d,** HPC terminal activation in the AHN (N=22). **e,f,** HPC terminal inhibition in the AHN (N=22). Viral spread in all injection areas (AHN, HPC) are depicted by the green blots across coronal plates of mouse brain atlas. Color bar, the number of mice with viral expression in the area (DAPI, blue; ChR2 and ArchT, green). Scale bar 200 µm.

**Extended Data Fig 2.**
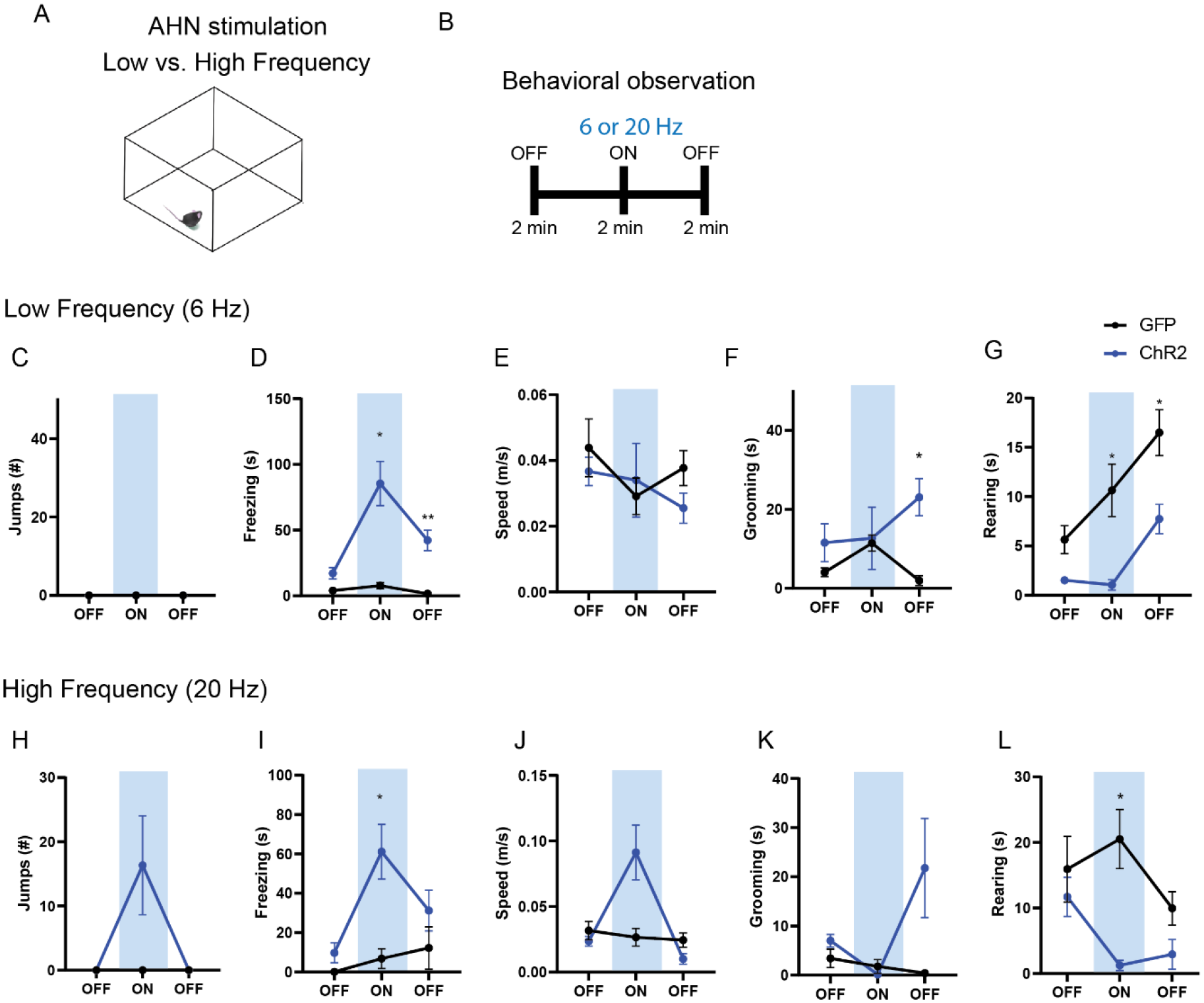
The effects of Low vs. High frequency AHN stimulation. (ChR2 N=6, GFP N=7). **a,** Schematic illustration of open field box where the low and high frequency stimulation occurred. **b,** Testing paradigm for with low vs. high frequency AHN stimulation. **c-g,** Behavioural changes induced by low frequency (6 Hz) AHN stimulation. **c,** No jumping observed upon light stimulation. **d,** Freezing was increased upon AHN stimulation (2-WAY RM ANOVA, light x genotype, F(2,22)=13.13, \*\*\**p=0.0002,* light effect, F (1.610, 17.71) = 16.95, *\*\*\*p=0.0002*, genotype effect, F (1, 11) = 35.33, *\*\*\*\*p<0.0001*, Sidak’s multiple comparison test, ON *\*p=0.0167,* Off 2 *\*\*p=0.0098*). **e,** Speed did not change (2-WAY RM ANOVA, light x genotype, F(2,22)=0.9463, *p=0.4034*, NS,), light effect, F (1.868, 20.54) = 1.426, *p=0.2622*, NS, genotype effect, F (1, 11) = 0.2379, *p=0.6352*, NS). **f,** Grooming was increased after AHN stimulation during the second light OFF period. (2-WAY RM ANOVA, light x genotype, F(2,22)=2.430, *p=0.1113,* NS, Sidak’s multiple comparison test, Off 2, *\*p=0.0164*). **g,** Rearing was decreased (2-WAY RM ANOVA, F(2,22)=2.407, *p=0.1135*, NS, F (1.691, 18.60) = 21.68, *\*\*\*\*p<0.0001*, F (1, 11) = 13.98, *\*\*p=0.0033*, Sidak’s multiple comparison test, ON, *\*p= 0.0317*, the second light OFF period, *\*p=0.0297*). **h-l,** Behavioural changes induced by high frequency (20 Hz) AHN stimulation. **h,** Jumping was increased upon AHN stimulation (2-WAY RM ANOVA, light x genotype, F (2, 22) = 5.328, *\*p=0.0130*, light effect, F (1.000, 11.00) = 5.328, *\*p=0.0414*, genotype effect, F (1, 11) = 5.328, *\*p=0.0414*). **i,** Freezing was increased upon AHN stimulation. (2-WAY RM ANOVA, light x genotype, F (2, 22) = 5.432, *\*p=0.0121*, light effect, F (1.772, 19.50) = 8.363, *\*\*p=0.0031*, genotype effect, F (1, 11) = 9.633, *\*p=0.01,* Sidak’s multiple comparison test, ON, *\*p= 0.0286*)**. j,** Speed was increased upon AHN stimulation, light x genotype, F (2, 22) = 15.55, *\*\*\*\*p<0.0001*, light effect, F (1.119, 12.30) = 15.12, *\*\*p=0.0017*, genotype effect, F (1, 11) = 2.073, *p=0.1777*, NS). **k,** Grooming was increased during the second light OFF period. (2-WAY RM ANOVA, light x genotype, F (2, 22) = 4.473, *p=0.0234, light effect, F (1.153, 12.69) = 3.175, p=0.0949, genotype effect, F (1, 11) = 6.429, *p=0.0277) **l,** Rearing is reduced upon AHN stimulation (2-WAY RM ANOVA, light x genotype, F (2, 22) = 5.564, *\*p=0.0111*, light effect, F (1.406, 15.47) = 4.790, *\*p=0.0339*, genotype effect, F (1, 11) = 6.128, *\*p=0.0308,* Sidak’s multiple comparison test, *\*p=0.0146*). All results reported are mean ± s.e.m. Sidak’s multiple comparison test: *p < 0.05, **p < 0.01, ***p < 0.001, ****p<0.0001.

**Extended Data Fig 3.**
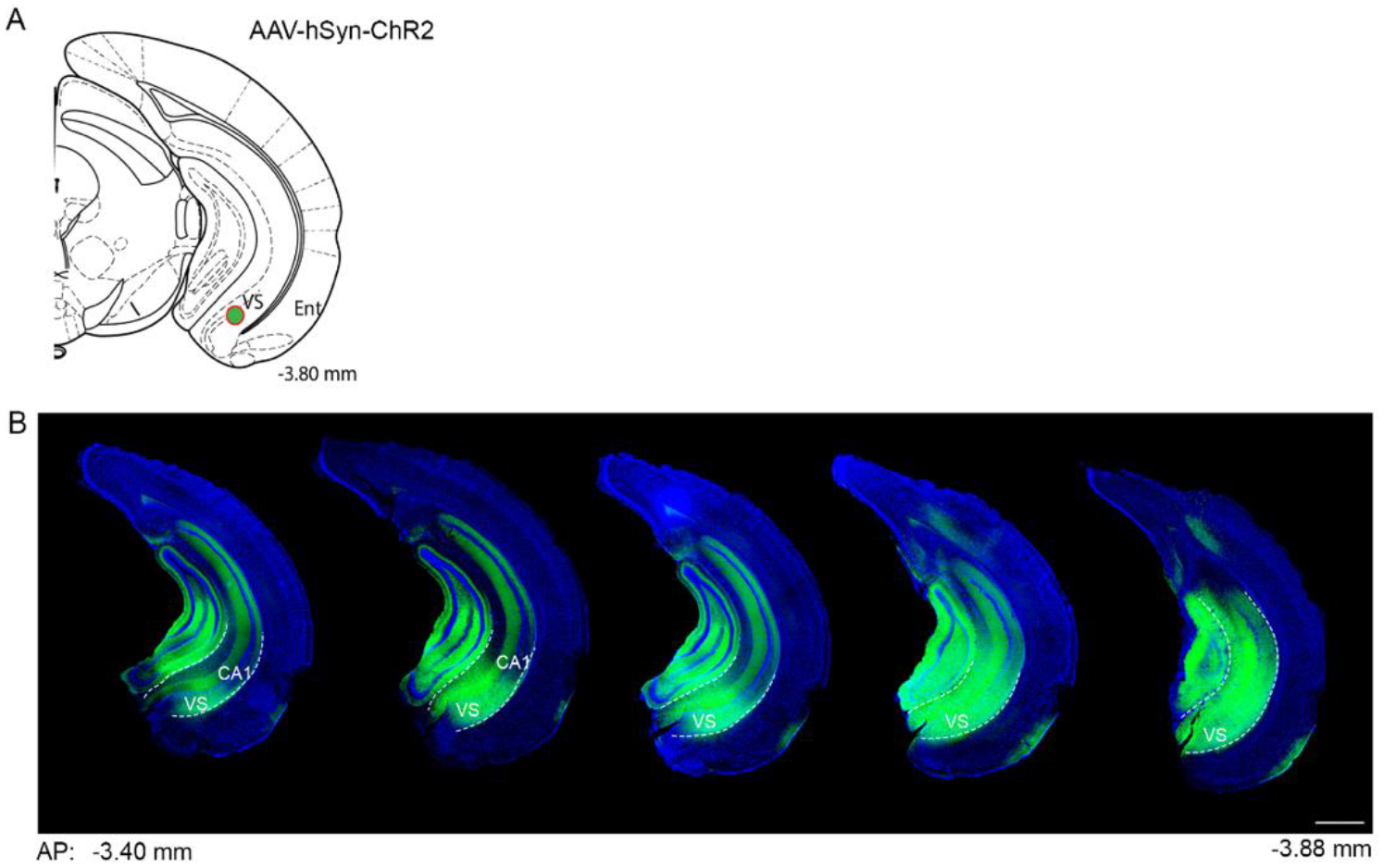
Viral expression of the anterograde tracer ChR2-GFP. **a,** Schematic showing the viral injection target (green dot) in the ventral subiculum. **b,** Representative coronal sections of the hippocampus showing the spread of AAV-hSyn-ChR2-GFP along the anterior-posterior axis of the hippocampus. DAPI, blue; ChR2, green; VS, ventral subiculum; CA1, cornu ammonis 1 of hippocampus. Scale bar = 500 µm

**Extended Data Fig 4.**
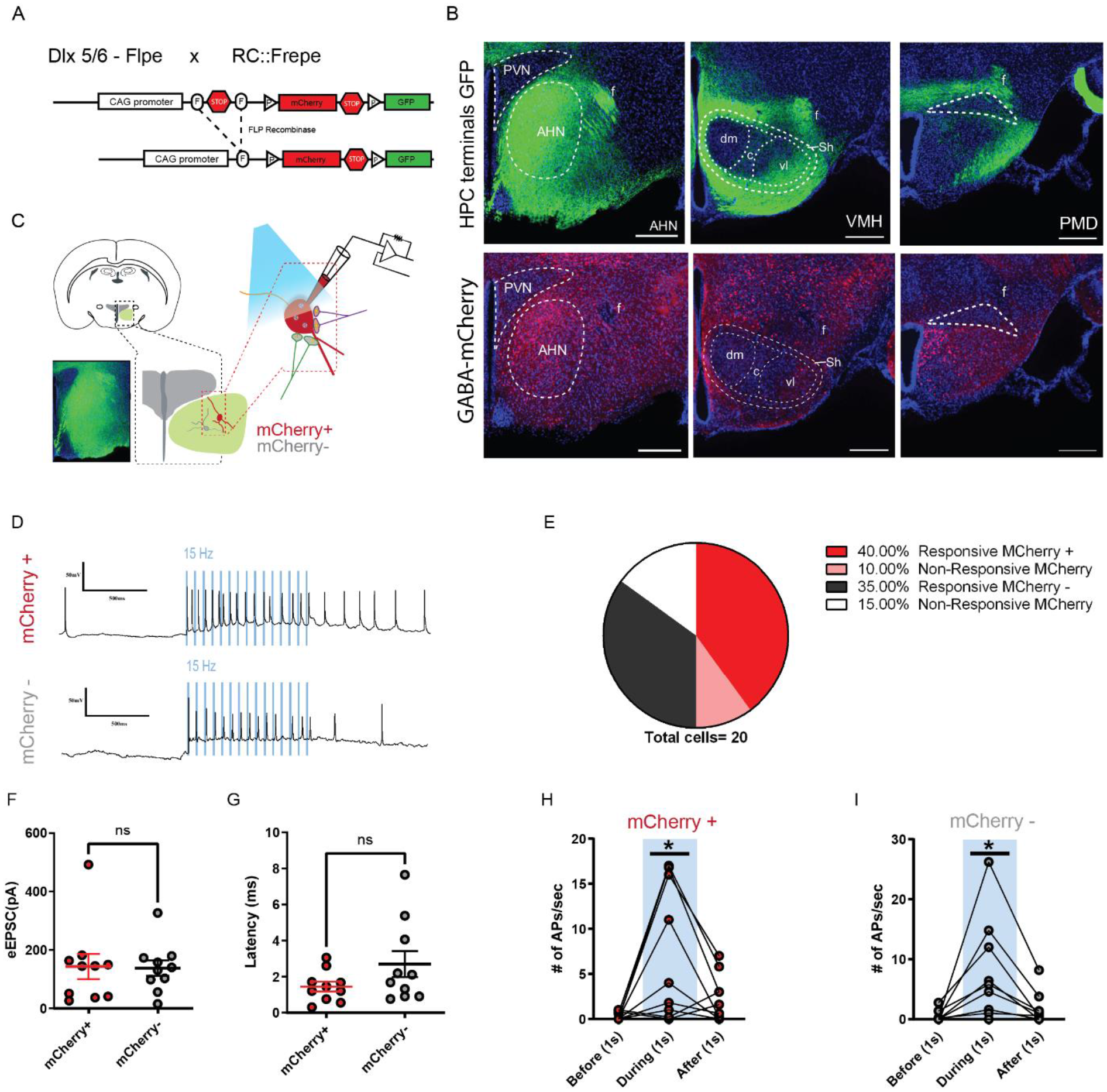
HPC inputs innervate both GABA and non-GABA cells in the AHN. **a,** Schematic showing the reporter allele, RC::FrePe, containing FRT-flanked and loxP-flanked transcriptional stop cassettes. Dlx 5/6 FLPe-mediated stop cassette removal results in mCherry expression in forebrain GABA cells. The RC::FrePe allele is knocked in to the Gt(ROSA)26Sor(R26) locus with CAG (chicken beta-actin and CMV enhancer) promoter elements. **b,** Top: HPC terminals (green) in the hypothalamus, including anterior hypothalamic nucleus (AHN), dorsomedial and central regions of ventromedial hypothalamus (VMHdm/c), premammillary dorsal nucleus (PMD), paraventricular nucleus (PVN), ventrolateral region of ventromedial hypothalamus (VMHvl), shell of ventromedial hypothalamus (VMHsh). DAPI staining (blue). Bottom: mCherry expression in GABA cells in the respective three regions from the same brain. **c,** Schematic illustration for patch clamp recordings of AHN neurons in coronal brain slices that express ChR2 in HPC terminals. Red: mCherry-positive GABA cells, Grey: mCherry-negative non-GABAergic cells. **d,** Examples of cell attached recordings. Illumination of blue light (480 nm, 5 ms pulse at 15 Hz) triggered action potential firing of AHN mCherry+ and mCherry-neurons. **e,** A pie chart depicting the percentage of AHN GABA or non-GABA cells that evoked light-evoked action potentials (mCherry-positive GABA cells, N=10; mCherry-negative non-GABAergic cells, N=10). **f,g,** Summary of light-evoked EPSC amplitude (**f**) and latency (**g**). **h,i,** Number of action potentials evoked 1s before, 1s during and 1s after light onset from mCherry-positive GABA cells (**h**) of the AHN (1-WAY RM ANOVA, F (1.119, 10.07) = 8.271, *\*p=0.0146*) and mCherry-negative non-GABAergic cells (**i**) of the AHN (1-WAY RM ANOVA, F (1.088, 9.791) = 7.116, \**p=0.0223*). All results reported are mean ± s.e.m. *\*p < 0.05*. Scale bar= 100µm

**Extended Data Fig 5.**
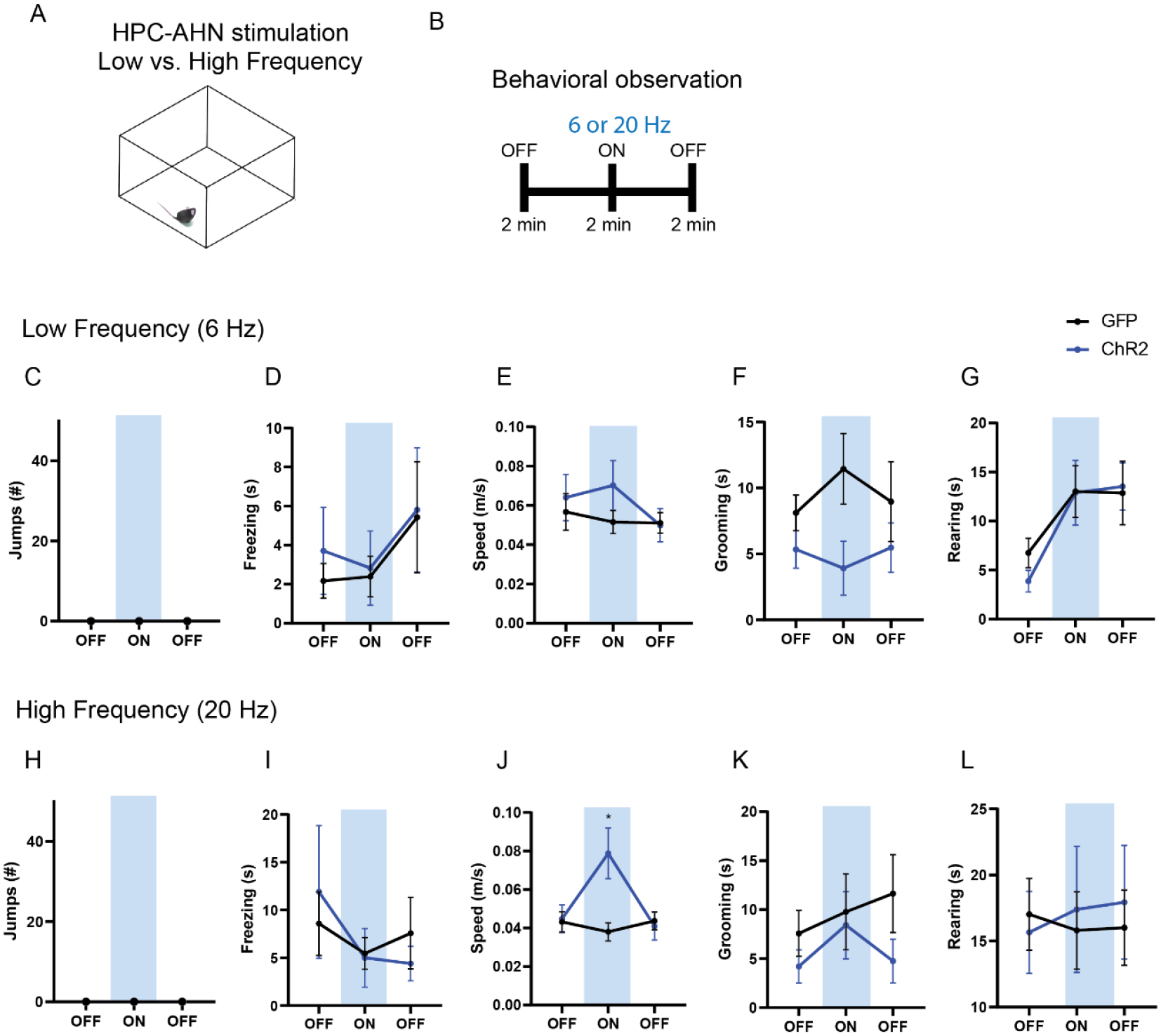
The effects of Low vs. High frequency HPC→AHN pathway activation. **a**, Schematic illustration of open field box where the low and high frequency HPC→AHN pathway stimulation occurred. **b**, Testing paradigm for with low vs. high frequency HPC→AHN pathway stimulation. **c-g,** Behavioural changes induced by low frequency (6 Hz) stimulation. **c,** No jumping observed upon stimulation. **d,** Freezing did not change (2-WAY RM ANOVA, light x genotype, F(2, 36) = 0.0762, *p=0.9268,* NS, light effect, F(1.040, 18.73) =1.947, *p=0.1793*, NS, genotype effect, F(1, 18)= 0.1065, *p=0.7479*, NS). **e,** Speed did not change (2-WAY RM ANOVA, light x genotype, F (2, 36) = 1.656, *p=0.2051*, NS, light effect, F(1.860, 33.48) = 2.281, *p=0.1211*, NS, genotype effect, F (1, 18) = 0.5189, *p=0.4806*, NS). **f,** Grooming was decreased upon stimulation (2-WAY RM ANOVA, light x genotype, F(2, 36) = 0.6101, *p=0.5488*, NS, light effect, F(1.640, 29.53) = 0.08480, *p=0.8846*, NS, genotype, F(1, 18) = 10.22, \*\**p=0.005*). **g,** Rearing did not change (2-WAY RM ANOVA, light x rhodopsin, F (2, 36) = 0.5721, *p=0.5694*, NS, light effect, F (1.927, 34.69) = 13.27, \*\*\*\**p<0.0001*, Sidak’s multiple comparison test, NS, rhodopsin effect, F (1, 18) = 0.07171, *p=0.7919*, NS). **h-l,** Behavioural changes induced by high frequency (20 Hz) stimulation. **h,** No jumping observed upon stimulation. **i,** Freezing did not change (2-WAY RM ANOVA, light x genotype, F (2, 36) = 0.5866, *p=0.5614*, NS, light effect, F(1.693, 30.47) = 1.614, *p=0.2173*, NS, genotype effect, F (1, 18) = 0.0006919, *p=0.9793*, NS). **j,** Speed was increased upon stimulation (2-WAY RM ANOVA, light x genotype, F (2, 36) = 15.26, \*\*\*\**p*<0.0001, light effect, F (1.779, 32.02) = 8.293, \*\**p*=0.0018, genotype effect, F (1, 18) = 1.916, *p=0.1832,* NS, Sidak’s multiple comparison test, ON, *\*p=0.0402*). **k,** Grooming did not change (2-WAY RM ANOVA, light x genotype, F(2, 36) = 0.3965, *p=0.6756,* NS, light effect, F (1.839, 33.09) = 0.5543, *p=0.5653,* NS, genotype effect, F(1, 18) = 2.738, *p=0.1153*.) **l,** Rearing did not change (2-WAY RM ANOVA, light x genotype, F (2, 36) = 0.2581, p=0.7739, NS, light effect, F (1.715, 30.86) = 0.03206, P=0.9524, NS, genotype effect, F (1, 18) = 0.03084, *p=0.8626*, NS). Sidak’s multiple comparison test : *\*p < 0.05, **p < 0.01, ***p < 0.001, ****p<0.0001*.

**Extended Data Fig 6.**
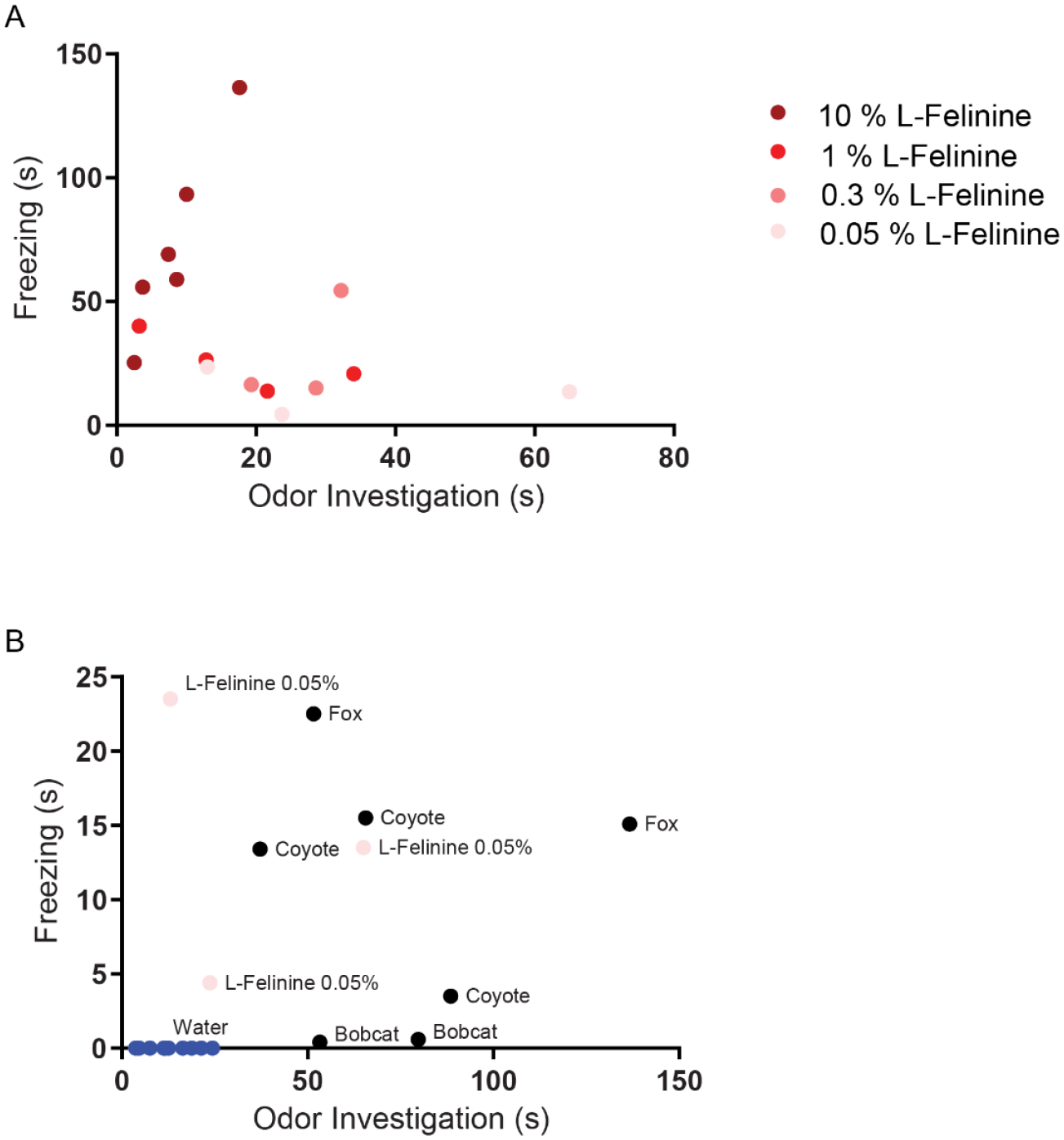
L-Felinine increases freezing in a dose-dependent manner. Each dot represents behavioural responses to different concentrations of L-Felinine that were measured in home cage during a 5 min trial. **a,** Freezing and odor investigation evoked by different concentrations of L-Felinine. Higher concentrations of L-Felinine induced greater freezing and shorter odor investigation. **b,** Freezing and odor investigation evoked by water, natural predator urine, and L-Felinine 0.05 % (concentration found in cat urine).

**Extended Data Fig 7.**
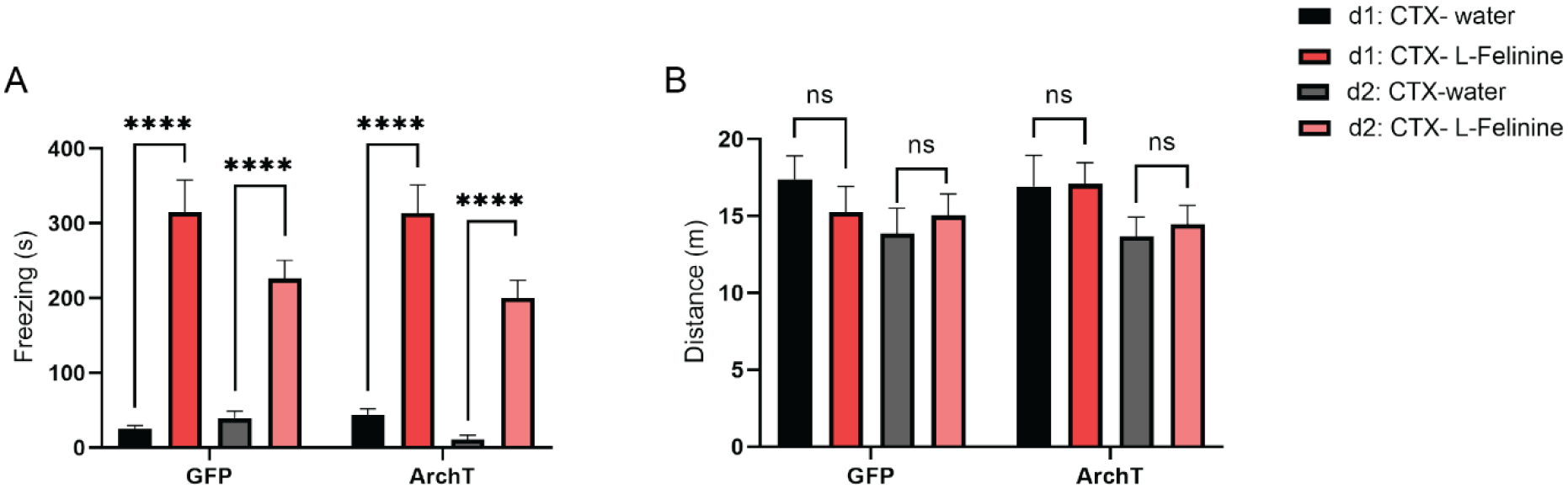
Behavioural responses to L-Felinine during conditioning sessions. **a**, Freezing induced by water and L-Felinine during two days of conditioning (d1 and d2) in GFP and ArchT mice (GFP N=8, ArchT N=12). Black: Day 1 water, Red: Day 1 L-Felinine, Grey: Day 2 water, Pink: Day 2 L-Felinine. (2-WAY ANOVA, D1 odorant x genotype, F(1,36)=0.2311, p=0.6332, D2 odorant x genotype, F(1,36)=0.2283, p=0.6353, \*\*\*\**p<0.0001*). b, Distance travelled during water and L-Felinine exposure during two days of conditioning (2-WAY ANOVA, D1 odorant x genotype, F(1,42)=0.4615,0.5007, 2-WAY ANOVA, D2 odorant x genotype, F(1,36)=0.01967, p=0.8891, NS). All results reported are mean ± s.e.m. Sidak’s multiple comparison test: *\*p < 0.05, **p < 0.01, ***p < 0.001, ****p<0.0001*.

**Extended Data Fig 8.**
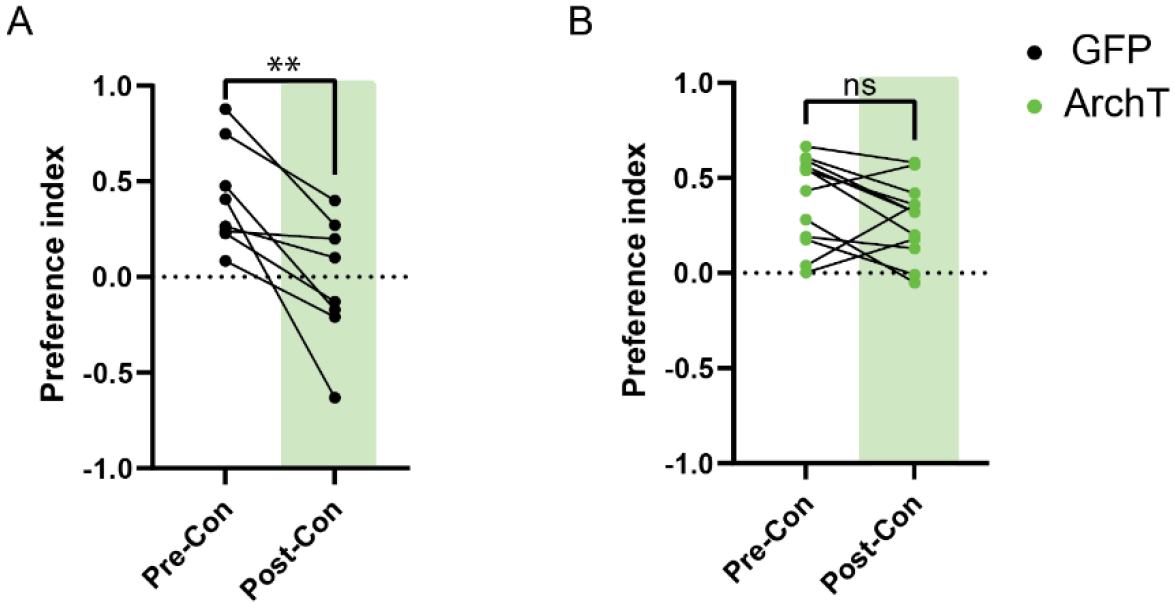
HPC→AHN pathway inhibition impairs the retrieval of place aversion memory. Preference index of GFP (**a**) and ArchT mice (**b**) (GFP = 8, ArchT N = 12) for the L-Felinine-coupled chamber during habituation (pre-con: pre-conditioning) and after conditioning session (post-con: post-conditioning). GFP pre-conditioning vs. GFP post-conditioning (paired t-test, t=3.381, df=7, *\*\*p=0.0096*). ArchT pre-conditioning vs. ArchT post-conditioning (paired t-test, t=1.704, df=11, p=0.1164).

**Extended Data Fig 9.**
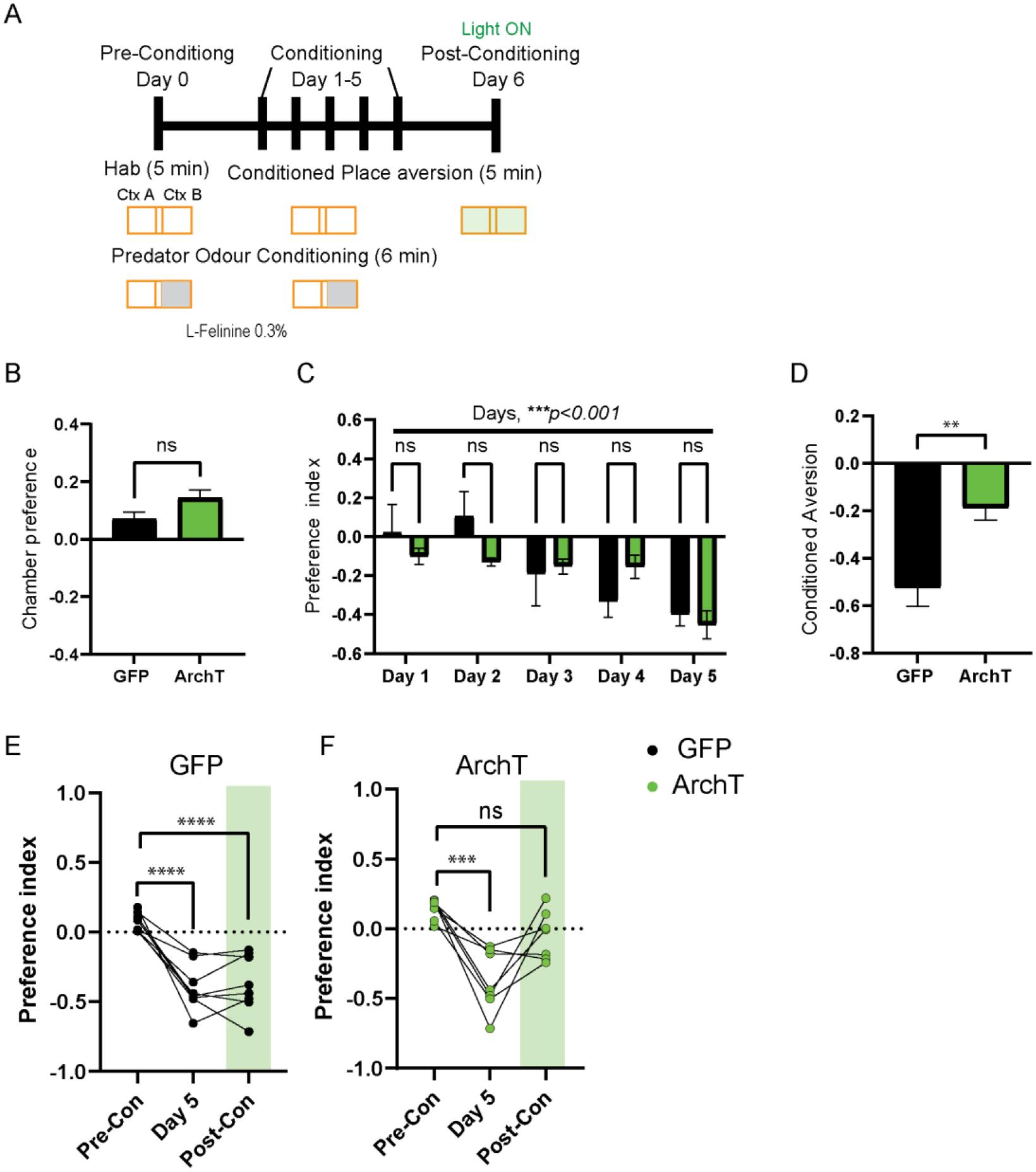
Development of predator associated context avoidance and impairment of retrieval of place aversion memory upon HPC-AHN pathway inhibition. **a,** Schematic illustration of the testing paradigm which consisted of multiple context-odor pairing sessions. It incorporated free chamber exploration (5 min) phase prior to the predator odor conditioning (6 min) with L-Felinine (0.3%). On Day 0, the free exploration or the Habituation (Hab), preferred chamber is selected as the predator odor chamber. **b,** Chamber preference during habituation (GFP N= 8, ChR2 N=7, unpaired t-test, t= 1.936, df= 13, *p=0.0749,* NS). **c**, Predator-context aversion developed across conditioning days. The day number equals the number of odour pairing throughout the training course. (2-WAY RM ANOVA, day x genotype, F(4,52)=1.53, *p=0.2071, NS,* day effect, F(2.923, 37.99)= 7.035, \*\*\**p=0.008*, genotype effect, F(1, 13)= 0.2547, *p=0.6222, NS*). **d,** Conditioned aversion memory. GFP vs. ArchT (unpaired t-test, t=3.514, df=13. *\*\*p=0.0038*). **e,** GFP animals developed conditioned aversion and light did not impair predator context retrieval (1-WAY RM ANOVA, Treatment effect, F(2,14)=31.51, *\*\*\*\*p<0.0001,* followed by Dunnett’s multiple comparison test for Pre-con vs. Day 5, *\*\*\*\*p<0.0001,* Pre-con vs. Post-Con, *\*\*\*\*p<0.0001).* **f**, ArchT animals developed conditioned aversion and light impaired predator context retrieval (1-WAY RM ANOVA, Treatment effect, F(2,12)=14.02, *\*\*\*p=0.0007,* followed by Dunnett’s multiple comparison test for Pre-Con vs. Day 5, *\*\*\*p=0.0004,* Pre-Con vs. Post-Con, *p=0.1358, NS*).

**Extended Data Fig 10.**
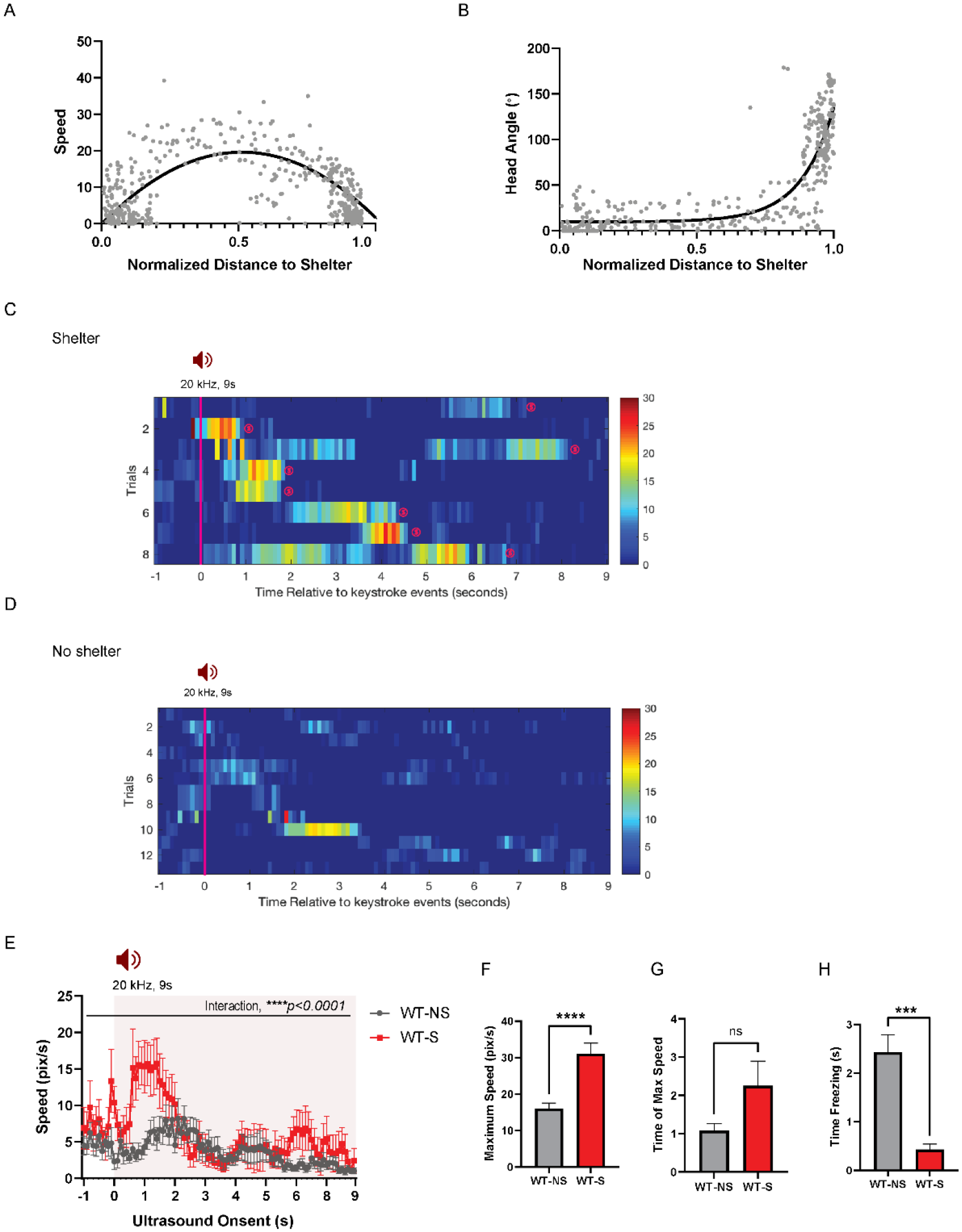
Mice display shelter-directed escape or freezing depending on the shelter availability. **a,** During ultrasound (US)-evoked escape responses, C57BL6 wild type mice (WT) run towards the shelter and reach the maximum speed in the middle of the escape trajectory to the shelter (WT-Best fit, quadratic, y=19.57 −3.757x+ 0.5372 - 75.48x^2^, maximum speed at x=0.5372). **b,** Mice turn their head toward the shelter quickly after they initiate escape flights (Best fit, sigmoidal, y= 7.433+ (1412478-7.433)/(1+10^Log0.4760-x^). **c,** Rastor plots of speed profile for WT animals’ escape flights to the shelter. **d,** Representative escape trajectories of wild type mice when shelter is not available. **Right,** Rastor plots of speed profile for WT animals’ escape flights when shelter is not available. Pink line denotes the US onset which lasted 9s. Animals’ arrivals at the shelter are denoted by “(s)”. Dot colors along the trajectory denote speed according to the color map on the right. **e,** Changes in speed (pix/s) 1 second before and 9 seconds after the US onset (2-WAY RM ANOVA, time x shelter availability, F(79,1981)=2.116, *\*\*\*\*p<0.0001*). **f,** Maximum speed during escape running (unpaired t-test, t=4.761, df=22, *\*\*\*\*p<0.0001*), **g,** Time elapsed to reach the maximum speed during escape running (unpaired t-test, t=1.869, df=22, p=0.075, NS). **h,** Time spent in freezing during the US presentation (unpaired t-test, t=4.266, df=19, *\*\*\*p=0.0004*). WT-NS: Wild type mice with no shelter. WT-S: Wild type mice with shelter. All results reported are mean ± s.e.m*. *p < 0.05, **p < 0.01, ***p < 0.001, ****p<0.0001*.

**Extended Data Fig 11.**
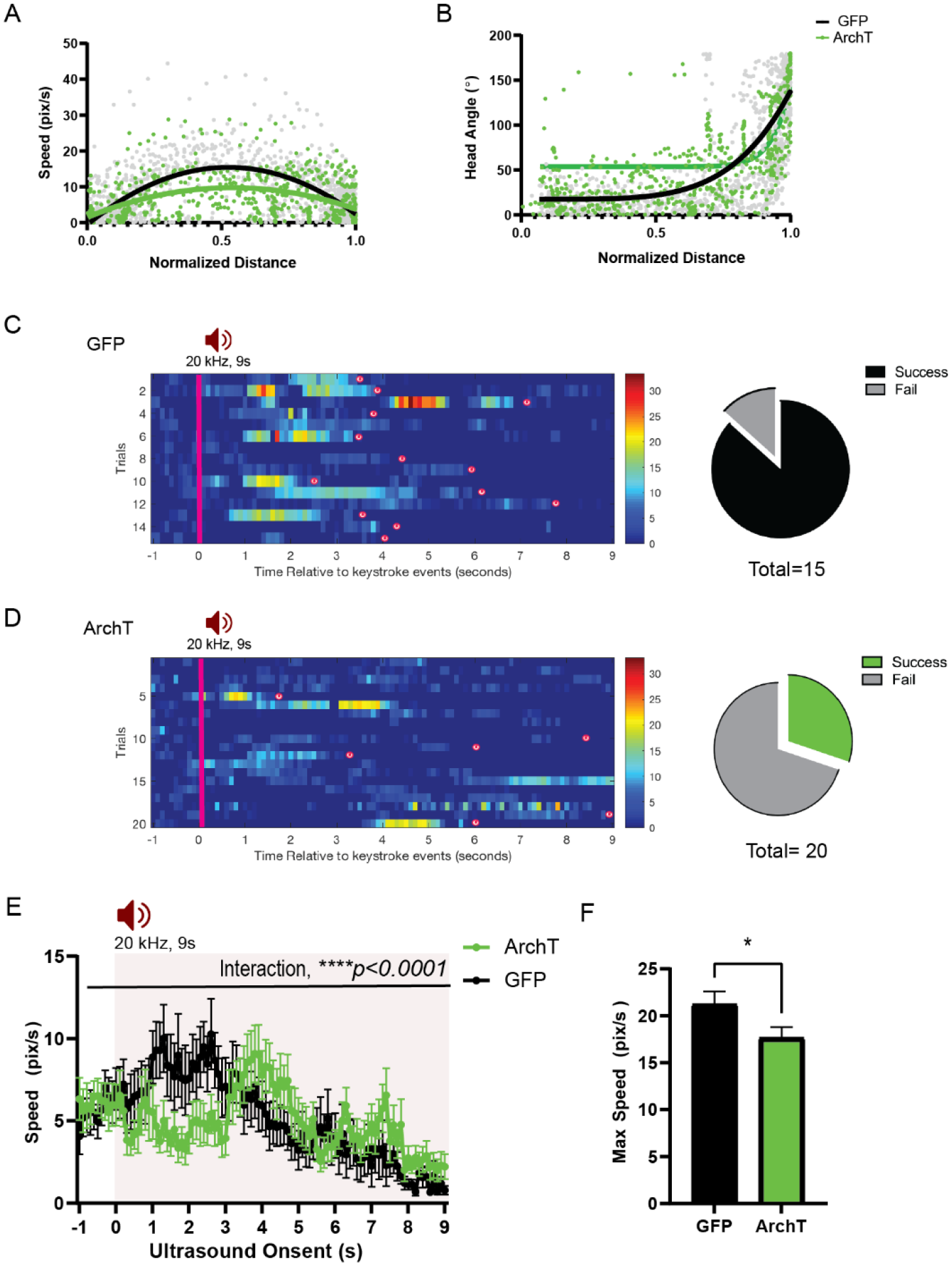
HPC→AHN pathway inhibition impairs ultrasound (US)-evoked escape responses. **a**, Comparison between GFP (N= 15) and ArchT (N=20) mice for speed (pixel/s) plotted in relation to the normalized distance to the shelter (GFP best fit line, y=15.34 −5.226x-58.01x^2^, maximum speed at x=0.5706; ArchT best fit line, y=9.768-0.2301x-28.23x^2^, maximum speed at x=0.5444). **b,** Head angle plotted in relation to the normalized distance to the shelter (GFP best fit line, y= 17.50+ (126075-4.765)/(1+10^Log0.6336-x^); ArchT best fit line, y=53.72+ (35041837-14.89)/(1+10^Log^0.3752^-x^)). **c, Left,** Rastor plots of speed profile for GFP. **Right,** Pie chart comparing the number of successful or failing escape responses upon ultrasound presentation (GFP: total 15 trials, 2 fails, 13 successes), **d, Left** Rastor plots of speed profile for ArchT. **Right,** Pie chart comparing the number of successful or failing escape responses upon ultrasound presentation (ArchT: total 20 trials, 14 fails, 6 successes). Pink line denotes the US onset which lasted 9s. Animals’ arrivals at the shelter are denoted by “(s)”. **e**, Changes in speed (pix/s) 1 second before and 9 seconds after the US onset (2-WAY RM ANOVA, time x genotype, F (100, 3300) = 1.763, \*\*\*\**p<0.0001*). **f**, Maximum speed during escape running (unpaired t-test, t=2.047, df=33, *\*p=0.0487*). All results reported are mean ± s.e.m. *p < 0.05, **p < 0.01, ***p < 0.001, ****p<0.0001.

**Extended Data Fig 12.**
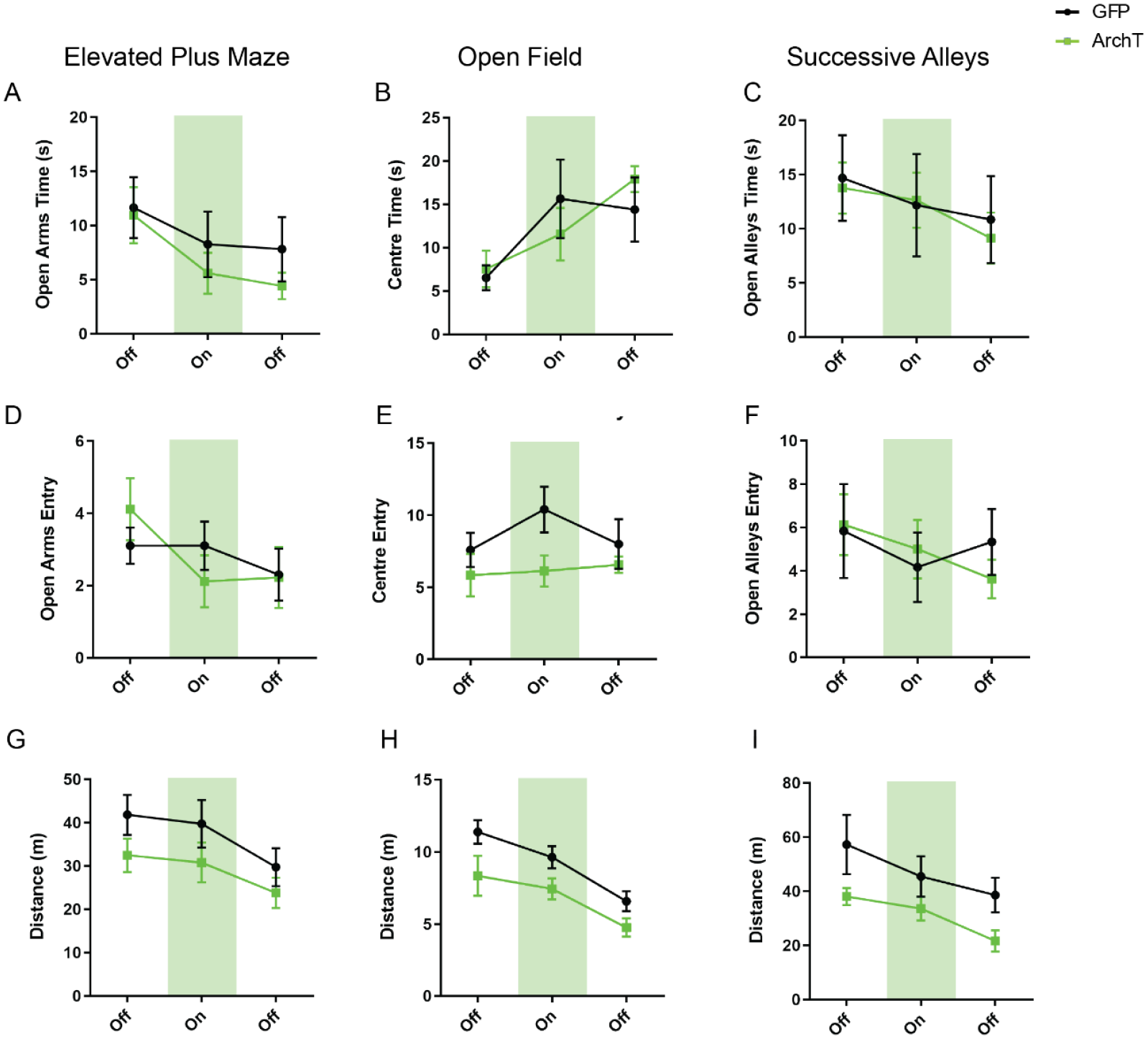
HPC→AHN pathway inhibition does not change anxiety-related behaviours. **a,** Time spent in the open arms in the elevated plus maze (EPM) (GFP N=10, ArchT N=9, 2-WAY RM ANOVA, time x genotype, F(2,34)=0.1412, NS). **b,** Time spent in the centre of the open field (OF) (GFP N=10, ArchT N=7, 2-WAY RM ANOVA, time x rhodopsin, F(2,30)=1.142, NS). **c,** Time spent in the open alleys of successive alleys (SA) (GFP N=6 ArchT N=8, 2-WAY RM ANOVA, time x genotype, F(2,24)=0.07654, NS). **d**, Number of entries into the open arms of EPM (2-WAY RM ANOVA, time x genotype, F(2,34)=1.578, NS). **e,** Number of entries into the centre of the OF (2-WAY RM ANOVA, time x rhodopsin F(2,30)=0.8540, NS). **f**, Number of entries into the open alleys of SA (2-WAY RM ANOVA, time x rhodopsin, F(2,24)=0.7369, NS). **g,** Distance travelled in the EPM (2-WAY RM ANOVA, F(2,34)=0.4192, NS). **h,** Distance travelled in the OF (2-WAY RM ANOVA, time x rhodopsin, F(2,30)=0.5938, NS). **i,** Distance travelled in the SA (2-WAY RM ANOVA, time x rhodopsin, F(2,24)=0.5972, NS). All results reported are mean ± s.e.m.

## References

1. Anderson, D. J. & Perona, P. Toward a science of computational ethology. Neuron 84, 18–31 (2014).

2. Gross, C. T. & Canteras, N. S. The many paths to fear. Nature Reviews Neuroscience 13, 651–658 (2012).

3. Silva, B. A., Gross, C. T. & Gräff, J. The neural circuits of innate fear: Detection, integration, action, and memorization. Learning and Memory 23, 544–555 (2016).

4. Evans, D. A., Stempel, A. V., Vale, R. & Branco, T. Cognitive Control of Escape Behaviour. Trends in Cognitive Sciences 23, 334–348 (2019).

5. Cooper, W. E. & Blumstein, D. Escaping from predators. An Integrative View of Escape Decisions (Cambridge University Press, Cambridge, 2015).

6. Vale, R., Evans, D. A. & Branco, T. Rapid Spatial Learning Controls Instinctive Defensive Behavior in Mice. Current Biology 27, 1342–1349 (2017).

7. Evans, D. A. Stempel, A.V., Vale, R., Ruehle, S., Lefler, Y. & Branco, T. A synaptic threshold mechanism for computing escape decisions. Nature 558, 590–594 (2018).

8. Blanchard, D. C. Risk assessment: at the interface of cognition and emotion. Current Opinion in Behavioral Sciences 24, 69–74 (2018).

9. Canteras, N. S. The medial hypothalamic defensive system: Hodological organization and functional implications. Pharmacology Biochemistry and Behavior 71, 481–491 (2002).

10. Fuchs, S. A. G., Edinger, H. M. & Siegel, A. The role of the anterior hypothalamus in affective defense behavior elicited from the ventromedial hypothalamus of the cat. Brain Research 330, 93–107 (1985).

11. Lamontagne, S. J., Olmstead, M. C. & Menard, J. L. The lateral septum and anterior hypothalamus act in tandem to regulate burying in the shock-probe test but not open-arm avoidance in the elevated plus-maze. Behavioural Brain Research 314, 16–20 (2016).

12. Wang, L., Chen, I. Z. & Lin, D. Collateral Pathways from the Ventromedial Hypothalamus Mediate Defensive Behaviors. Neuron 85, 1344–1358 (2015).

13. Silva, B. A. Mattucci, C., Kryzwkowski P., Cuozzo, R., Carbonari, L. & Gross, C.T. The ventromedial hypothalamus mediates predator fear memory. European Journal of Neuroscience 43, 1431–1439 (2016).

14. Pérez-Gómez, A. et al.Innate predator odor aversion driven by parallel olfactory subsystems that converge in the ventromedial hypothalamus. Current Biology 25, 1340–1346 (2015).

15. Canteras, N. S. Chapter 3.2 Neural systems activated in response to predators and partial predator stimuli. Handbook of Behavioral Neuroscience (2008).

16. Dielenberg, R. A., Hunt, G. E. & McGregor, I. S. “When a rat smells a cat”: The distribution of Fos immunoreactivity in rat brain following exposure to a predatory odor. Neuroscience 104, 1085–1097 (2001).

17. Blanchard, D. C., Canteras, N. S., Markham, C. M., Pentkowski, N. S. & Blanchard, R. J. Lesions of structures showing FOS expression to cat presentation: Effects on responsivity to a Cat, Cat odor, and nonpredator threat. Neuroscience and Biobehavioral Reviews 29, 1243– 1253 (2005).

18. Martinez, R. C. R., Carvalho-Netto, E. F., Amaral, V. C. S., Nunes-de-Souza, R. L. & Canteras, N. S. Investigation of the hypothalamic defensive system in the mouse. Behavioural Brain Research 192, 185–190 (2008).

19. Mendes-Gomes, J. et al. Defensive behaviors and brain regional activation changes in rats confronting a snake. Behavioural Brain Research 381, 112469 (2020).

20. Canteras, N. S. & Goto, M. Fos-like immunoreactivity in the periaqueductal gray of rats exposed to a natural predator. NeuroReport 10, (1999).

21. Tovote, P. et al. Midbrain circuits for defensive behaviour. Nature 534, 206–212 (2016).

22. Lammers J.H,. Kruk M.R., Meelis W., & van der Poel A.M. Hypothalamic substrates for brain stimulation-induced patterns of locomotion and escape jumps in the rat. Brain Res. 449, 294–310 (1988)

23. Siegel, A. & Pott C.B. Neural substrate of aggression and flight in the cat. Progress in Neurobiology 31, 261–283 (1988).

24. Lipp, H. P., & Hunsperger, R. W. Threat, attack and flight elicited by electrical stimulation of the ventromedial hypothalamus of the marmoset monkey Callithrix jacchus. Brain, behavior and evolution, 15, 260–293 (1978).

25. Wilent, W. B. et al. Induction of panic attack by stimulation of the ventromedial hypothalamus. Journal of neurosurgery, 112(6), 1295–1298 (2010).

26. Maren, S., Phan, K. L. & Liberzon, I. The contextual brain: Implications for fear conditioning, extinction and psychopathology. Nature Reviews Neuroscience 14, 417–428 (2013).

27. Pentkowski, N. S., Blanchard, D. C., Lever, C., Litvin, Y. & Blanchard, R. J. Effects of lesions to the dorsal and ventral hippocampus on defensive behaviors in rats. European Journal of Neuroscience 23, 2185–2196 (2006).

28. Kjelstrup, K. G. et al. Reduced fear expression after lesions of the ventral hippocampus. Proceedings of the National Academy of Sciences of the United States of America 99, 10825– 10830 (2002).

29. Wang, M. E. et al. Differential roles of the dorsal and ventral hippocampus in predator odor contextual fear conditioning. Hippocampus 23, 451–466 (2013).

30. Lisman, J., Buzsáki, G., Eichenbaum, H. et al. Viewpoints: How the hippocampus contributes to memory, navigation and cognition. Nature Neuroscience 20, 1434–1447 (2017).

31. Aqrabawi, A. J. & Kim, J. C. Hippocampal projections to the anterior olfactory nucleus differentially convey spatiotemporal information during episodic odour memory. Nature Communications 9, 2735 (2018).

32. Mandairon, N. et al. Context-driven activation of odor representations in the absence of olfactory stimuli in the olfactory bulb and piriform cortex. Frontiers in Behavioral Neuroscience 8, 138, (2014).

33. Gottfried, J. A., Smith, A. P. R., Rugg, M. D. & Dolan, R. J. Remembrance of odors past: Human olfactory cortex in cross-modal recognition memory. Neuron 42, 687–695 (2004).

34. Clinchy, M., Sheriff, M. J. & Zanette, L. Y. Predator-induced stress and the ecology of fear. Functional Ecology 27, 56–65 (2013).

35. Takahashi, L. K., Nakashima, B. R., Hong, H. & Watanabe, K. The smell of danger: A behavioral and neural analysis of predator odor-induced fear. Neuroscience and Biobehavioral Reviews 29, 1157–1167 (2005).

36. Canteras, N. S. & Swanson, L. W. Projections of the ventral subiculum to the amygdala, septum, and hypothalamus: A PHAL anterograde tract-tracing study in the rat. Journal of Comparative Neurology (1992)

37. Risold, P. Y. & Swanson, L. W. Connections of the rat lateral septal complex. Brain Research Reviews 24, 115–195 (1997).

38. Parfitt, G. M. et al. Bidirectional Control of Anxiety-Related Behaviors in Mice: Role of Inputs Arising from the Ventral Hippocampus to the Lateral Septum and Medial Prefrontal Cortex. Neuropsychopharmacology 42, 1715–1728 (2017).

39. Amaral, D. G. & Witter, M. P. The three-dimensional organization of the hippocampal formation: A review of anatomical data. Neuroscience 31, 571–591 (1989).

40. Bienkowski, M. S. et al. (2018). Integration of gene expression and brain-wide connectivity reveals the multiscale organization of mouse hippocampal networks. Nature neuroscience, 21, 1628–1643.

41. Friedman, D. P., Aggleton, J. P. & Saunders, R. C. Comparison of hippocampal, amygdala, and perirhinal projections to the nucleus accumbens: Combined anterograde and retrograde tracing study in the macaque brain. Journal of Comparative Neurology 450, 345–365 (2002).

42. Hoover, W. B. & Vertes, R. P. Anatomical analysis of afferent projections to the medial prefrontal cortex in the rat. Brain Structure and Function 212, 149–179 (2007).

43. Petreanu, L., Mao, T., Sternson, S. M., & Svoboda, K. (2009). The subcellular organization of neocortical excitatory connections. Nature, 457(7233), 1142–1145.

44. Voznessenskaya, V.V., Kvasha, I. G., Klinov, A. B. & Laktionova, T. K. Responses to Domestic Cat Chemical Signals in the House Mouse Are Modulated by Early Olfactory Experience. in Chemical Signals in Vertebrates 13, 401–411 (2016).

45. Hendriks, W. H., Moughan, P. J., Tarttelin, M. F., & Woolhouse, A. D. Felinine: a urinary amino acid of Felidae. Comparative biochemistry and physiology. Part B, Biochemistry & molecular biology, 112, 581–588 (1995).

46. Blanchard, D. C., Agullana, R. & Squirrels, G. Twenty-two kHz alarm cries to presentation of a predator, by laboratory rats living in visible burrow systems. Physiology and Behavior 50, 967–972 (1993).

47. Litvin, Y., Blanchard, D. C. & Blanchard, R. J. Vocalization as a social signal in defensive behavior. Handbook of Behavioral Neuroscience, 19, 151–157 (2010).

48. Maren, S. & Quirk, G. J. Neuronal signalling of fear memory. Nature Reviews Neuroscience 5, 844–852 (2004).

49. Anagnostaras, S. G., Gale, G. D. & Fanselow, M. S. Hippocampus and contextual fear conditioning: Recent controversies and advances. Hippocampus 11, 8–17 (2001).

50. Boudaba, C., Szabó, K. & Tasker, J. G. Physiological mapping of local inhibitory inputs to the hypothalamic paraventricular nucleus. Journal of Neuroscience 16, 7161–7160 (1996).

51. Ziegler, D., Cullinan W. & Herman, J. Distribution of vesicular glutatmate transporter mRNA in rat hypothalamus. *Journal of Comp*. Neurology 448, 217–229 (2002).

52. Kishi, T., Tsumori, T., Ono, K., Yokota, S., Ishino, H., & Yasui, Y. Topographical organization of projections from the subiculum to the hypothalamus in the rat. Journal of Comparative Neurology 419, 205–222 (2000).

53. Cezario, A. F., Ribeiro-Barbosa, E. R., Baldo, M. V., & Canteras, N. S. Hypothalamic sites responding to predator threats - The role of the dorsal premammillary nucleus in unconditioned and conditioned antipredatory defensive behavior. European Journal of Neuroscience 28, 1003–1015 (2008).

54. Nguyen, R. et al. Cholecystokinin-Expressing Interneurons of the Medial Prefrontal Cortex Mediate Working Memory Retrieval. The Journal of neuroscience 40, 2314–2331 (2020)

55. Taniguchi, H. et al. A Resource of Cre Driver Lines for Genetic Targeting of GABAergic Neurons in Cerebral Cortex. Neuron, 71, 995–1013 (2011).

56. Whissell, P. D. & Bang, J. Y. et al. Selective activation of cholecystokinin-expressing GABA (CCK-GABA) neurons enhances memory and cognition. eNeuro 6, 0360–18 (2019).

57. Dedic, N., Kühne, C., Jakovcevski, M., Hartmann, J., Genewsky, A. J., Gomes, K. S., Anderzhanova, E., Pöhlmann, M. L., Chang, S., Kolarz, A., Vogl, A. M., Dine, J., Metzger, M. W., Schmid, B., Almada, R. C., Ressler, K. J., Wotjak, C. T., Grinevich, V., Chen, A., Schmidt, M. V., … Deussing, J. M. Chronic CRH depletion from GABAergic, long-range projection neurons in the extended amygdala reduces dopamine release and increases anxiety. Nature Neuroscience 21, 803–807 (2018).

58. Bang, S. J., Jensen, P., Dymecki, S. M. & Commons, K. G. Projections and interconnections of genetically defined serotonin neurons in mice. European Journal of Neuroscience 35, 85–96 (2012).

59. McNaughton, B. L., Battaglia, F. P., Jensen, O., Moser, E. I., & Moser, M. B. Path integration and the neural basis of the ‘cognitive map’. Nature reviews. Neuroscience, 7(8), 663–678 (2006).

60. Whishaw, I. Q., & Gorny, B. Path integration absent in scent-tracking fimbria-fornix rats: evidence for hippocampal involvement in “sense of direction” and “sense of distance” using self-movement cues. The Journal of neuroscience 19, 4662–4673 (1999).

61. Xia, L., Nygard, S. K., Sobczak, G. G., Hourguettes, N. J., & Bruchas, M. R. Dorsal-CA1 Hippocampal Neuronal Ensembles Encode Nicotine-Reward Contextual Associations. Cell reports, 19, 2143–2156 (2017).

62. Hsiang, H. L. et al. Manipulating a “cocaine engram” in mice. The Journal of neuroscience 34, 14115–14127 (2014).

